# Voltage imaging reveals that hippocampal interneurons tune memory-encoding pyramidal sequences

**DOI:** 10.1101/2023.04.25.538286

**Authors:** Jiannis Taxidis, Blake Madruga, Maxwell D Melin, Michael Z Lin, Peyman Golshani

## Abstract

Hippocampal spiking sequences encode and link behavioral information across time. How inhibition sculpts these sequences remains unknown. We performed longitudinal voltage imaging of CA1 parvalbumin- and somatostatin-expressing interneurons in mice during an odor-cued working memory task, before and after training. During this task, pyramidal odor-specific sequences encode the cue throughout a delay period. In contrast, most interneurons encoded odor delivery, but not odor identity, nor delay time. Population inhibition was stable across days, with constant field turnover, though some cells retained odor-responses for days. At odor onset, a brief, synchronous burst of parvalbumin cells was followed by widespread membrane hyperpolarization and then rebound theta-paced spiking, synchronized across cells. Two-photon calcium imaging revealed that most pyramidal cells were suppressed throughout the odor. Positive pyramidal odor-responses coincided with interneuronal rebound spiking; otherwise, they had weak odor-selectivity. Therefore, inhibition increases the signal-to-noise ratio of cue representations, which is crucial for entraining downstream targets.

## INTRODUCTION

The hippocampus is critical for transforming a sequence of experiences into memories while also keeping track of their temporal relationships [1]. Sequentially activated pyramidal ensembles are thought to underlie this function [1, 2]. Some hippocampal pyramidal cells respond to the perception of a sensory cue to be remembered (*‘cue cells*’), generating sensory representations [3, 4, 5, 6]. Other pyramidal cells (‘*time cells’)*, fire in succession after the cue, encoding its memory as well as the time elapsed since its presentation [5, 7, 8, 9, 10]. Spiking sequences of cue cells followed by cue-specific time cells, provide a neural mechanism for linking contiguous experiences across gaps in time into a memory trace [2, 11, 12].

Population activity patterns, including such spiking sequences, require the close coordination between excitation and inhibition in neuronal networks [13]. Yet the role of inhibitory neurons in sculpting memory-encoding hippocampal dynamics is largely unknown. Inhibition in the hippocampus is regulated by a diverse array of GABAergic interneurons, divided into subtypes with distinct anatomical organization, connectivity, firing patterns and genetic profiles [14, 15]. Two major subtypes are parvalbumin-expressing (PV) and somatostatin-expressing (SST) interneurons which mostly target perisomatic areas and dendrites of pyramidal cells, respectively. Perisomatic PV-inhibition controls the timing of pyramidal output [16] and coordinates rhythmic population activity [17, 18, 19]. Dendritic SST-inhibition gates the integration of proximal and distal dendritic inputs on pyramidal cells [20] and ultimately their gain (their output relative to their input) [21]. The interplay between pyramidal cells, PV and SST interneurons shapes hippocampal spiking *in vivo* and controls behavior (reviewed in [22, 23, 24, 25]). However, this interplay has rarely been studied outside of spatial navigation. As a result, the role of PV and SST neurons in sculpting memory-encoding spiking sequences, combining cue cells and time cells, remains unknown.

Exploring the role of inhibition in memory processes requires monitoring activity from identified interneuron subtypes with high temporal resolution, to reliably record changes in their high firing rates. It also requires tracking the same cells across memory-relevant timescales of multiple days. Electrophysiology does not allow combined cell-type specificity and multi-day cell tracking, whereas calcium imaging cannot capture the fast-spiking dynamics of interneurons. Moreover, neither calcium imaging nor extracellular electrophysiology captures subthreshold membrane potentials which contain valuable information on synaptic inputs. Fast frame-rate voltage imaging of genetically encoded voltage indicators (GEVIs) [26, 27] overcomes these barriers and captures action potentials as well as subthreshold membrane potentials. It also allows imaging of the same cells for multiple days [28].

We employed kHz-rate voltage imaging to investigate the spiking and membrane potential dynamics of interneurons. PV and SST cells from mouse dorsal CA1 were recorded with the ASAP3 GEVI [28], while mice were passively exposed to odors or actively performing an odor-cued working memory behavior [29]. Some cells were tracked for multiple days, as well as before and after task training. We previously described pyramidal sequences during this task, composed of ‘odor cells’, encoding specific odor cues, followed by odor-specific time cells, encoding time-points in the delay period after a specific odor cue [5]. Here, we found that unlike pyramidal cells, many PV and SST cells increased spiking during the presentation of the odor cue but were not selective to any odor. In addition, very few cells had time fields during the delay period. Interestingly, interneurons could retain the same odor field before and after training. On a population level, odor-timed inhibition was driven by a stable number of interneurons daily, with a constant turnover, and was not altered by task learning. At odor onset, a highly synchronous, transient spiking of PV cells, but not SST cells was followed by widespread hyperpolarization across cells. This hyperpolarization reset intracellular theta oscillations and organized rebound odor-spiking into synchronous theta cycles. We assessed the effect of this widespread, synchronized, theta-modulated inhibition, during odor presentation, on pyramidal sequences using two-photon calcium imaging of CA1 pyramidal cells during the same task. We found that (1) most pyramidal cells were silenced during odor cues, (2) the spiking of the few odor-encoding cells was coincident with the interneuronal rebound spiking otherwise (3) if pyramidal spiking occurred earlier, during interneuronal hyperpolarization, it showed weak odor selectivity. Therefore, PV and SST inhibition suppressed background pyramidal spiking, temporally tuned odor-encoding responses, and enhanced odor selectivity. Collectively inhibition increased the signal-to-noise ratio of excitatory cue cells. This sculpting of sensory representations is crucial for reliably transferring memory-relevant information to cortical targets downstream.

## RESULTS

### *In vivo* voltage imaging reveals PV-cell and SST-cell spiking and subthreshold dynamics during the DNMS working memory task

To capture the fast-scale spiking and subthreshold membrane potential dynamics of PV and SST interneurons, we performed voltage imaging *in vivo*.

We expressed the ASAP3 GEVI in hippocampal PV or SST interneurons, by virally transfecting the right dorsal CA1 region of adult PV-Cre and SST-Cre transgenic mice (N = 5 mice for each group) with Cre-dependent ‘ASAP3’ virus (AAV8-ef1α-DiO-ASAP3-Kv) [28]. Mice were implanted with an imaging window above the corpus callosum and their skull was affixed with a metal headbar for head-fixation (Figure 1A). Mice were later water-deprived and trained to perform an olfactory delayed non-match-to-sample (DNMS) task, requiring working memory, while head-fixed on a spherical treadmill (Figure 1A) [5, 29]. An odor cue was presented for 1 sec and, after a 5 sec delay, it was followed by a second odor. Mice were trained to lick a lickport to release water rewards only if the two odors did not match and to refrain from licking if the odors matched (Figure 1B). Odor cues were either isoamyl acetate (‘odor A’) or pinene (‘odor B’) and were delivered in random combinations. Odor-match trials with licking were not punished, but water rewards were not delivered. We have previously shown that, after a shaping period, mice learn to perform this task within ~5-6 days using working memory [5]. Performance was quantified as the ratio of summed correct hits and correct rejections over all trials.

**Figure 1:**
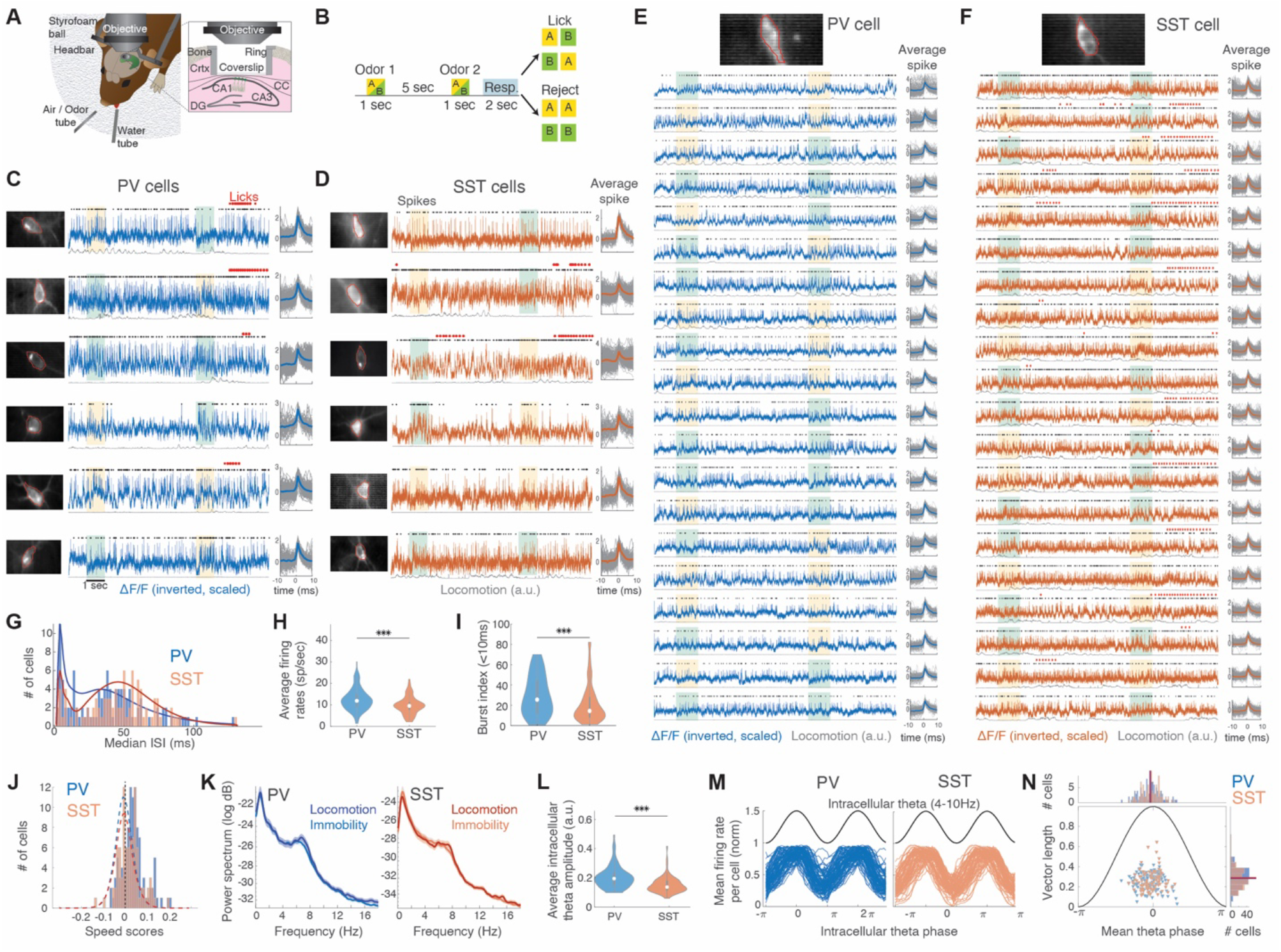
*In vivo* voltage imaging with ASAP3 of CA1 PV and SST interneurons during DNMS trials. **A**. Illustration of the behavioral and experimental set-up. Crtx: Cortex. CC: corpus callosum. **B**. Schematic of the DNMS trial. ‘Odor-A’: isoamyl acetate (yellow). ‘Odor-B’: pinene (green). Licking is assessed during a 2 sec response window (blue). **C-D**. Example traces from PV (**C**) and SST (**D**) interneurons during a DNMS trial (from N = 3 and 5 mice respectively). Left: Motion corrected mean field of view with the ROI outlined (red). Middle: Inverted ΔF/F, scaled by maximum value. Black dots: Detected spikes. Red: Licks. Color boxes indicate the two odors in the trial as in **B**. Gray traces: Locomotion. Right: ΔF/F segments during all individual (gray) and average (thick) action potentials in the trial. **E-F**. Example PV (**E**) and SST (**F**) interneuron recorded during 20 continuous DNMS trials, depicted as in **C-D. G**. Histogram of median inter-spike interval per cell and estimated probability density (solid lines). **H-I**. Average firing rates (**H**) and burst index (**I**) for PV vs SST interneurons. *** P < 0.001, Wilcoxon ‘ranksum’ test (WT). **J**. Histogram of average speed score per cell. Dashed lines: Mean shuffle baseline per cell group. **K**. Mean ± SE power spectral density of each cell group during concatenated motion versus immobility segments. No significant differences exist at any frequency (P > 0.05; WT; FDR). **L**. Average intracellular theta amplitude (4-10 Hz) per cell for PV vs SST interneurons. *** P < 0.001; WT. **M**. Mean firing rate of each PV (left) and SST interneuron (right), normalized to their maximum value, plotted as function of the intracellular theta cycle (black). **N**. Mean phase locking strength (vector length) versus preferred theta phase for each cell during the intracellular theta cycle (triangles). Top: Histogram of preferred phase for the two cell groups. Right. Same for vector length. Lines indicate distribution means. No significant differences observed (Parametric Watson-Williams multi-sample test and WT respectively). White circles in violin plots indicate median values throughout.

We performed voltage imaging of PV or SST interneurons for two imaging sessions (two separate days) while mice were naïve to the task and passively smelled the odor cues. We then trained mice to perform the DNMS task (~10-14 days total) and resumed voltage imaging after mice were well-trained to the task (see Methods). Imaging took place with a custom-made epifluorescence microscope at 1000 frames per second (1 kHz). The field of view (FOV) was 88×44 µm (64×32 pixels), which in most cases encompassed a single cell. Recordings captured action potentials and subthreshold membrane potential fluctuations (Figure 1C-D). We were able to record activity across multiple trials (up to 48 from a single cell), retaining clear spike waveforms and subthreshold dynamics, without photo-bleaching cells (Figures 1E-F). We recorded 107 PV cells and 93 SST cells in total (in 104 and 90 videos respectively), with an average of 16.2 ± 6.5 trials per PV cell (mean ± SD) and 14.7 ± 8.5 trials per SST cell.

PV cells fired with lower interspike intervals than SST cells (Figure 1G), yielding higher average firing rates (Figure 1H) and burst indices (Figure 1I) [16]. Spiking was mostly modulated by the DNMS task, rather than by locomotion on the treadmill, as evidenced by the overall low speed-scores (see Methods; 79.4% PV and 85% SST cells had scores below chance levels; Figure 1J) as well as by the similar spiking during motion and immobility in PV cells (though SST rates were higher during motion; Figure S1). Power spectra of ΔF/F traces, after action potential waveform removal (see Methods), contained a peak in theta frequency range (4-10 Hz) with similar power during motion and immobility (Figure 1K). Average intracellular theta amplitude was significantly higher for PV cells than SST cells (Figure 1L) and was correlated with average firing rates only in PV cells (Figure 1S). Firing rates were strongly phase-locked to the intracellular theta peak, with similar phase locking strength between PV and SST cells (Figures 1M-N).

Collectively, *in vivo* fast-rate voltage imaging, with the ASAP3 GEVI, reliably captured spiking and membrane potential dynamics of PV and SST interneurons during the DNMS task.

### Most PV and SST interneurons encode odor delivery, but not odor identity or delay-time

During the DNMS task, CA1 pyramidal cells form odor-specific spiking sequences of ‘odor cells’, whose activation encoded the presentation of a specific odor, followed by ‘time cells’, each spiking at a particular timepoint in the delay after a specific odor [5]. Here, we asked if interneurons form similar odor-specific spiking sequences.

Since pyramidal sequences were also observed in untrained mice, we pooled all sessions, including those when the animal was naïve to the DNMS task. We focused our analysis on the period covering the first odor presentation and the delay period in each trial (‘odor-delay interval’). Interneurons generally fired continuously, but many cells increased their firing rates during odor presentation (Figures 2A,D). To assess the existence of odor-related or temporal firing fields, we detected significant peaks in each cell’s average firing rate (binned at 100 msec) throughout the odor-delay interval and computed the cell’s odor selectivity index (see Methods).

**Figure 2:**
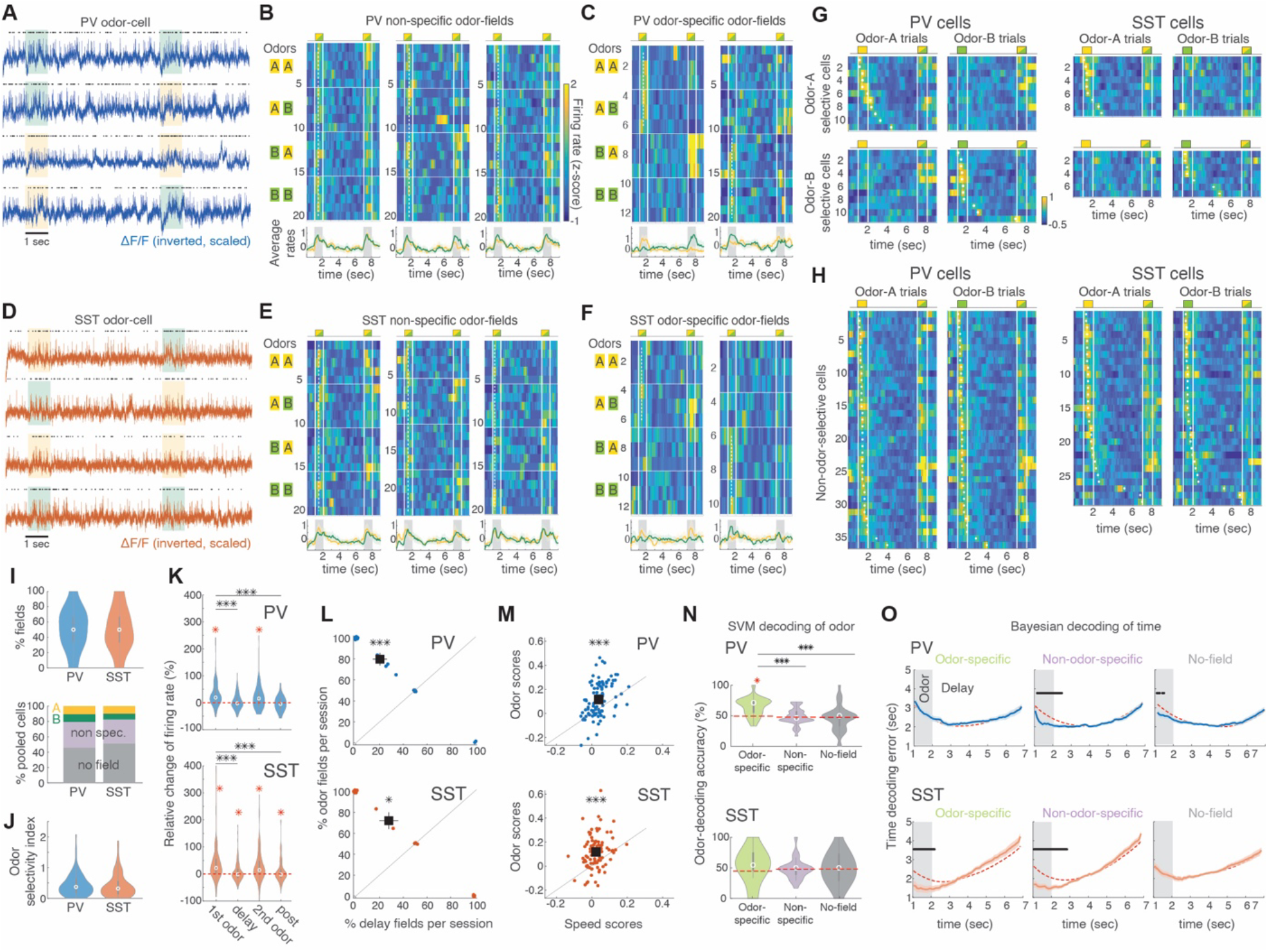
PV and SST interneurons encode the presentation of the odor-cue, not delay-time, with weak odor-selectivity. **A**. Example traces from a PV odor field across 4 trials, displayed similarly to Fig 1. Spiking increases during the presentation of either odor cue. **B**. Example PV odor fields, encoding both odor-A and odor-B presentation (left panel is same cell as in **A**). Each row is the neuron’s z-scored firing rate during a trial, with trials stacked in blocks (horizontal lines) according to the odor combination (shown in left boxes). Vertical lines: odor delivery (trial layout shown on top). Dashed lines: firing field time-bin. Bottom: Mean ± SE rate over all odor-A (yellow) and odor-B trials (green). **C**. Example odor-specific PV interneurons significantly encoding either odor-A (left) or odor-B (right), plotted as in **B** with same color-scale. Dashed lines cover only the trials corresponding to the preferred odor. **D-F**. Same as **A-C** but for SST interneurons. **G**. Left 4 panels: Average z-scored firing rates of odor-A-specific (top row) and odor-B-specific (bottom row) PV cells over odor-A (left) and odor-B (right) trials, stacked by their field time-bin (white dots). The second odor in the trial can be either one. Right 4 panels: Same for SST interneurons. **H**. Left 2 panels: Average firing rates of all non-odor-specific PV cells over odor-A (left) and odor-B (right) trials, stacked by their field time-bin. Right 2 panels: Same for SST cells with same color-scale. **I**. Top: Percentage of PV and SST field cells per mouse per session. Bottom: Mean cumulative ratio of odor-specific cells, non-odor-specific ones, and cells with no field. No significant differences observed between PV-SST (P > 0.05, WT). **J**. Odor-selectivity (absolute values) of pooled cells between the two cell groups (P > 0.05, WT). **K**. Relative change in average firing rates, compared to baseline, during the first odor, the delay, the second odor, and the post-odor response window in PV and SST field cells. No differences exist between the first and second odor. Distributions with means significantly higher than zero indicated by red asterisk (P < 0.05, right-tailed t-test, FDR-corrected). *** P < 0.001, WT. **L**. Percentage of odor fields versus delay time fields over all field cells in each session, in PV and SST cells. Small jitter added for clarity. Black square: Distribution means. * P < 0.05; *** P < 0.001, paired-sample t-test. **M**. Odor-scores vs speed scores for PV and SST cells. Black square: Distribution means. *** P < 0.001, paired-sample t-test. **N**. Odor-decoding accuracy with SVM decoders trained only on odor-specific cells, non-odor-specific cells, or no-field cells, during odor-presentation. *** P < 0.001, WT, FDR-corrected. Dashed lines: Mean shuffle baselines. Only odor-specific PV cells performed better than chance (red * P < 0.001, WT, FDR-corrected). **O**. Average time-decoding error (absolute values) as function of trial time, with Bayesian decoders trained on each cell group as in **N**. Dashed lines: Mean shuffle baselines. Black bars: Time bins with error significantly lower than chance. P < 0.05 WT, FDR-corrected.

Many interneurons exhibited firing fields during the odor presentation but these were independent of the odor’s identity (Figures 2B,E). Only a few cells had odor-specific fields (Figure 2C,F), yielding sparse odor-specific sequences when pooled across mice and sessions (11 PV and 8 SST cells were odor-A specific; 10.3% and 8.6% of all cells in each group. 11 PV and 7 SST cells were odor-B specific; 10.3% and 7.5% respectively; Figure 2G). In contrast, non-odor-specific fields were found in 36 PV and 30 SST cells (33.6% and 33.2% of all cells in each group; Figure 2H). The percentage of detected fields per session was similar between PV and SST interneurons, with similar ratios of odor-A- and odor-B-specific and non-odor-specific fields (Figure 2I). The average odor selectivity was also similar between PV and SST cells (Figure 2J). In total, 54.2% of all PV cells and 48.4% of SST cells had a significant field within the odor-delay interval (termed ‘field cells’), whereas the remaining 45.8% and 51.6% cells showed no significant fields (‘no-field cells’).

Firing rates of PV and SST field cells increased similarly during the first and second odor in a trial and were significantly higher compared to the delay period and the response window (Figure 2K). During the delay, PV firing rates returned to baseline levels whereas SST firing rates remained higher than baseline (Figure 2K). As a result, most fields were during odor delivery (‘odor fields’). During the delay period interneuronal firing rates were rarely time-modulated, resulting in very sparse ‘delay fields’ compared to odor fields (Figure 2L; 9.3% of PV and 12.9% of SST cells had fields during the delay, with 50% and 58.3% of those, respectively, having their field within 1 sec from odor offset).

Locomotion did not drive spiking during the odor delivery. This was assessed by computing odor scores similarly to speed scores for all cells (see Methods). 55.1% of all PV cells and 58.1% SST cells had significant odor scores, compared to 20.6% of PV cells and 15% of SST cells for speed scores. As a result, odor scores were significantly higher than speed scores for PV and SST field-cells (Figure 2M).

Furthermore, odor responses were not triggered by clicking sounds by the odor-delivery air-valve. In a set of recordings, the odor airflow was turned off on alternate trials while all valves remained operational. PV and SST spiking was reduced during odor OFF trials compared to odor ON trials. This reduction was often seen throughout the odor-delay interval (Figure S2). On average PV and SST cells exhibited significantly lower firing during OFF trials, even though their average locomotion was similar in the two conditions (Figure S2).

To further assess whether interneuronal responses contained information on odor identity, we trained binary support vector machine (SVM) classifiers on each field cell’s activity over the first odor. As expected, SVM classifiers trained on odor-specific PV cells, decoded odor identity better than chance, and better than those trained on non-odor-specific cells or no-field cells, both of which performed at chance levels (Figure 2N). All SST-trained classifiers performed at chance levels, even those trained on odor-specific cells (Figure 2N). Classifiers trained on the entire odor-delay interval, instead of just the first odor cue, performed at chance levels (P > 0.05 WT; compared to shuffle baseline).

We also trained Bayesian decoders on each cell’s firing rate to decoded time throughout the odor-delay interval. The prevalence of odor fields and lack of delay fields resulted in better than chance time-decoding during the odor delivery but chance levels during the delay period for most PV and SST cell groups (Figure 2O). Odor decoding through the Bayesian decoders was at chance levels in all cases (P > 0.05 compared to shuffle baseline for all bins).

Collectively, our findings reveal widespread PV and SST inhibition during odor delivery, independently of odor identity, and sparse time-locked inhibition during the delay. These characteristics are unlike the highly odor-selective spiking of pyramidal cell sequences that tile the entire delay period.

### Population inhibition is in ‘steady state’ across days and independent of DNMS training

Pyramidal spiking sequences combine stable odor fields across days with highly dynamic delay time fields. Delay fields are also learning-dependent, increasing in number as mice learn the DNMS task [5]. We thus examined whether similar stability and learning-related dynamics hold for interneuronal firing fields.

We assessed changes in inhibition across days or between naïve and trained states by tracking a subset of neurons across multiple imaging sessions. To ensure a cell was the same as one recorded previously, we recorded a few frames from each neuron with an extended FOV (352×352 µm at < 100 Hz frame rate). The average frame was used as reference to make accurate comparisons based on processes and other features surrounding the cell body. We were able to perform voltage imaging from the same interneuron for up to 5 consecutive imaging sessions (different days, Figure S3) while the mouse performed the DNMS task, as well as record cells before and after DNMS training (Figures 3A,C).

**Figure 3:**
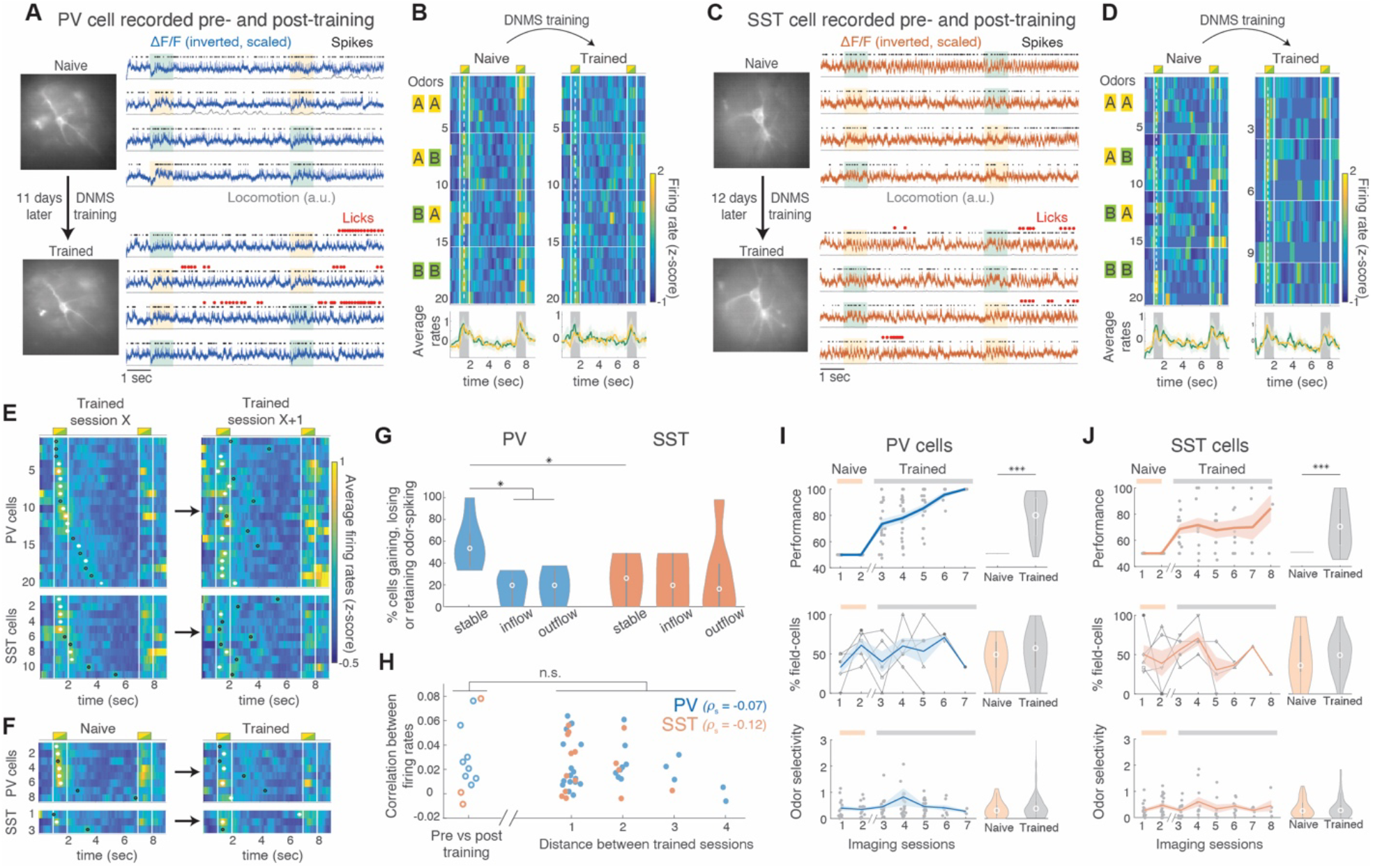
Interneuronal stability dynamics across days and before versus after DNMS training. **A**. Example PV cell recorded first during passive DNMS exposure and after DNMS training. Left: expanded field of view to display entire cell and surroundings. Right: ΔF/F, detected spikes, locomotion and licking during 4 DNMS trials. **B**. Firing rate per trial and odor-specific average rate across the two recording sessions, plotted as in Fig 2. **C-D**. Same as **A-B** for an example SST cell. Both cells retained a significant non-specific odor field across days and DNMS training. **E**. Average firing rate across all trials for pooled PV (top) and SST cells (bottom) on any recording session X and the following one, X+1. Cells are stacked according to their maximum rate time-bin on session X in both cases. White dots: Significant fields. Black circles: non-significant firing peak time-bins. **F**. Same as **E** for cells recorded before and after DNMS training (last naïve session vs first trained session). **G**. Percentage of PV and SST cells recorded across two consecutive sessions that retained odor-spiking (‘stable’), moved their firing peak into (‘inflow’) or out of the odor delivery time bins (‘outflow’). * P < 0.05; WT; FDR. **H**. Mean cross-correlation of a cell’s firing rates across all trials between two sessions, as a function of the distance between the sessions. Pre-Post: Same for correlations between last naïve vs first trained session (P > 0.05, WT; between the two groups for both cell types). Small jitter added for clarity. *ρ*_S_: Spearman correlation between distance of trained sessions and firing rate correlations (P > 0.05 for both cell groups). **I**. Top row: Performance of pooled PV-Cre mice across trials per cell (dots) and mean± SE over all trials per day (lines). Naïve and trained sessions indicated on top (no licking in naïve sessions results in 50% performance score). Note that multiple days of training take place between the two session groups. Middle row: Same for percentage of field cells over all recorded cells per session. Markers and lines indicate individual mice. Bottom row: Same for odor-selectivity of PV-cells (absolute values). Right panels: Corresponding distributions for naïve versus trained sessions pooled; only significant differences indicated (*** P < 0.001, WT). **J**. Same for SST-Cre mice and SST cells accordingly.

In general, fields tended to remap across days. The average firing rate peak of a cell could become a significant field or lose significance, shift along the time axis, or switch between being odor-specific or non-specific (Figure S3). As a result, odor selectivity indices fluctuated around zero across days (Figure S3).

Surprisingly, some PV an SST cells exhibited the same odor field in a naïve and in a trained session, weeks apart (Figure 3B,D). Moreover, PV odor fields were often retained between a trained session X and the next session X+1, whereas most PV delay fields turned to odor fields (Figure 3E). Similarly, 3 out of the 5 recorded PV odor fields in a naïve session remained odor fields in the first trained session (Figure 3F). SST fields exhibited higher instability overall (Figures 3E,F).

We quantified the turnover of odor fields across days by counting the percentage of cells from any session X+1 that retained an odor field from session X (‘stable’) or gained one (‘inflow’) as well as the percentage of cells in session X that lost their odor field in session X+1 (‘outflow’). There were significantly more stable PV odor fields than their inflow or outflow. Stable PV fields were also significantly more than stable SST fields, and the latter did not differ from inflow or outflow rates (Figure 3G).

We assessed a cell’s spiking similarity between any two sessions, by computing the average correlation of its firing rates between all pairs of trials of the two sessions. Correlations were low and they did not differ between trained sessions compared to before versus after training (Figure 3H). Moreover, there was no relationship between correlations and the distance between imaging sessions (Figure 3H). These findings corroborate an overall random change in firing activity between days, which was not affected by learning the DNMS task.

We also searched for any learning-related changes in the collective activity of all recorded cells. Unlike the increase in pyramidal delay fields throughout learning, we found stable percentages of PV and SST field cells across days, as well as pre- versus post-training, even though performance improved across trained sessions (Figures 3I-J). The average PV and SST odor selectivity also remained stable (Figures 3I-J), corroborating our observation in the multiday-tracked cells.

Finally, even though locomotion on the treadmill increased significantly after mice were trained, compared to naïve sessions, the firing rates of multiday-tracked PV and SST cells did not change over days, and only the average rates across all SST cells showed a small but significant increase after training (Figure S3). Intracellular theta power and theta phase locking of spikes increased after training only in PV cells. An increase in theta power was also observed in multiday-tracked cells when pooling PV and SST cells together, though it remained stable across trained sessions (Figure S3).

Collectively, these findings demonstrate a steady state inhibition by a stable number of cells over days, with fluctuating spiking modulation, irrespective of naïve exposure or participation to the DNMS task. Interestingly, PV cells had a higher probability of retaining odor fields for days and even across training.

### A hyperpolarization of PV and SST interneurons at odor onset resets intracellular theta oscillations

Little is known about subthreshold dynamics of interneurons *in vivo*, which contain valuable information on behaviorally relevant synaptic inputs. We next focused on the subthreshold membrane potential signals from PV and SST interneurons.

A hyperpolarizing negative deflection was prominent in the membrane potential of PV and SST interneurons during odor onset (Figure 4A,C). This was observed in multiple cells and across multiple trials and was reflected in a cell’s average de-spiked ΔF/F across both odor-A and odor-B trials (Figure 4B,D). Similar negative deflections could be seen during the delay but did not occur in a repeated manner (Figure 4A,C). The sharp hyperpolarization was generally followed by a depolarization during later stages of the odor delivery (during odor spiking), together yielding a delta-frequency wave (0.5-3 Hz) in the membrane potential (Figure 4B,D).

**Figure 4:**
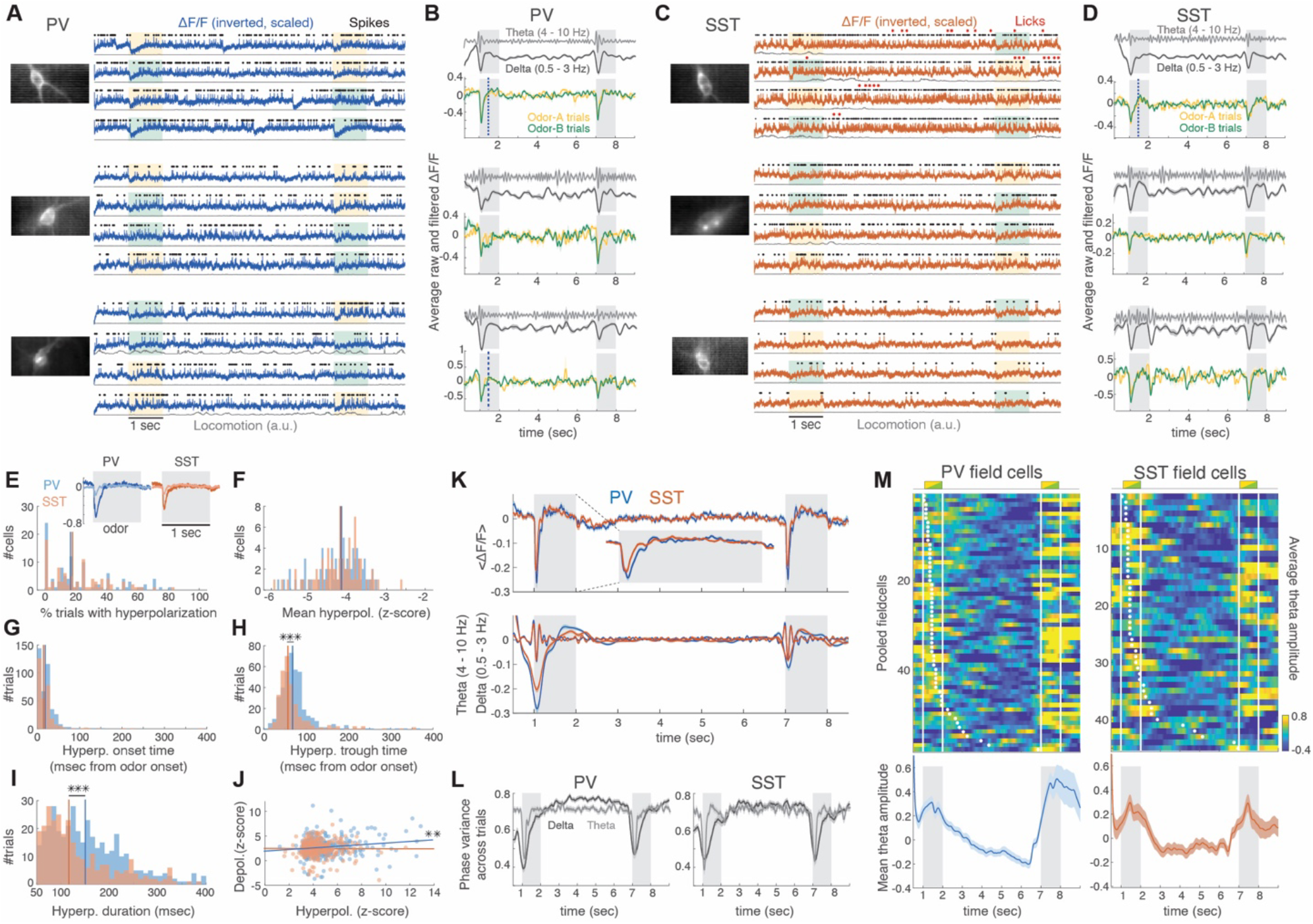
A hyperpolarization during odor-onset resets intracellular theta. **A**. Example trials from 3 PV cells (from 3 different mice) with odor-triggered hyperpolarization traces. Left: average field of view. Right: ΔF/F, detected spikes and locomotion during 4 DNMS trials, displayed as before. **B**. Odor-A (yellow) and Odor-B (green) average ΔF/F across all corresponding trials, for the 3 cells shown in **A**. Top panels: Trial-average delta-filtered (blue) and theta-filtered ΔF/F (gray). **C-D**. Same as **A-B** for 3 example SST cells (from 3 different mice). **E**. Percentage of trials with a significant hyperpolarization per cell. Lines indicate median values. Insets: Average odor-triggered ΔF/F over trials with significant and non-significant hyperpolarization. **F**. Distribution of average amplitude of ΔF/F hyperpolarization per cell (z-score-scaled over baseline of 0.5 sec before odor onset). **G-I**. Distribution of hyperpolarization onset time (**G**), time of minimum ΔF/F (**H**) and hyperpolarization duration (**I**). Distributions truncated at 400 msec for clarity. *** P < 0.001 WT; FDR. **J**. Amplitude of maximum depolarized versus hyperpolarized ΔF/F during odor-presentation. Lines: Least square fit for each cell type. Both correlations are highly significant. P* P < 0.01, F-test, for both cell types. One outlier removed for plotting clarity. **K**. Trial-average ΔF/F (top) and delta- and theta-frequency bandpassed ΔF/F (bottom) for PV (blue) and SST cells (red). Inset: Zoomed in segment around the first odor. **L**. Variance of delta and theta phases across all trials for each cell group. **M**. Average intracellular theta power across all trials for PV (left) and SST (right) field cells. Cells stacked by their time field (dots). Bottom: Theta power averaged across all field cells.

Significant hyperpolarizations (see Methods) occurred in a similar percentage of trials in PV and SST cells (22.5%± 21.5% versus 21.7%± 18.9%; P > 0.05, WT) and were absent in 22.4% PV and 19.3% SST cells (Figure 4E). During significant hyperpolarizations, the average, de-spiked ΔF/F deflections were similar between PV and SST cells (−4.8±1.7 versus −4.4±1.2 standard deviations of the baseline before the odor; P > 0.05, WT; Figure 4F). Hyperpolarization onset times were also similar (20±20 msec versus 23±60 msec after odor-onset; P > 0.05, WT; Figure 4G). PV hyperpolarizations peaked slightly but significantly later than SST ones (81±70 msec versus 83±97 msec post odor-onset – due to more outliers in SST cells – but with medians at 68msec versus 57 msec; P < 0.05; WT, FDR; Figure 4H) and lasted significantly longer (194±166 msec versus 187±203 msec post odor-onset; P < 0.05; WT, FDR; Figure 4I). Moreover, the amplitude of significant hyperpolarizations was correlated with the ensuing depolarizations during the odor in PV cells only (Figure 4J) suggesting a relationship between this initial input and the ensuing odor field spiking.

The fact that this negative deflection was not stereotypically reproduced in all cells and trials argues against it being artifactual. However, to ensure it was not driven by vibration or motion artifacts, we detached the odor- and air-delivery valve from the recording rig (this valve was the only moving hardware during odor onset). The same hyperpolarizations were found before and after valve detachment (Figure S4). Furthermore, to ensure the hyperpolarization was not driven by any artifactual increase in fluorescence (since ΔF/F is inverted) of the negatively tuned ASAP3, we expressed the positively tuned ASAP4 GEVI [30] in a set of mice (N = 5 SST-IRES-Cre mice). Non-inverted ASAP4 ΔF/F traces yielded similar negative deflections as ASAP3, corroborating that these deflections correspond to membrane hyperpolarization rather than artifacts.

To test if hyperpolarizations were linked to odor detection by the mouse or a different behavioral variable, we compared ΔF/F traces when the odors were turned OFF compared to ON. We observed similar traces in both conditions (Figure S4), suggesting that odor detection was not driving the inhibitory synaptic input. Furthermore, hyperpolarization amplitudes were similar during the delivery of preferred and non-preferred odors in odor-specific field cells, indicating they do not arise from specific odor cues (Figure S4). Hyperpolarization amplitudes and the frequency of significant hyperpolarizations across trials were also similar between odor-specific cells, non-odor-specific ones, and no-field cells (Figure S4).

Interestingly, the hyperpolarization amplitude in PV cells dropped significantly after training compared to naïve sessions, whereas it did not change for SST cells (Figure S4). The subsequent depolarizations during the odor did not change after training for either cell group (Figure S4).

The mean duration of the hyperpolarization places it in the theta frequency range (~5 Hz). Indeed, theta bandpassed traces, averaged across trials, showed an increase (Figures 4B,D), nested within the delta wave of the hyperpolarization-followed-by-depolarization cycle during the odor delivery (Figure 4K). This increase in average bandpassed ΔF/F indicates a transient theta phase resetting, rather than a transient increase in theta amplitude. This was supported by a reduction in theta phase variance across trials at odor onset (Figure 4L). Theta amplitudes exhibited a broader increase around a cell’s field, not just at odor-onset (Figure 4M), in accordance with previous observations of a similar increase of theta amplitude around a place cell’s spatial field [31].

Collectively, these findings reveal a novel inhibitory input on PV and SST interneurons triggered by the onset of odor delivery, but dissociated from odor detection, that transiently resets intracellular theta oscillations.

### Odor-onset hyperpolarization is preceded by brief synchronous PV output and synchronizes rebound spiking at theta cycles

We next examined the spiking correlates of PV and SST interneurons to odor-onset hyperpolarization on a finer timescale.

PV cells often produced a single spike or short burst, immediately preceding the ensuing hyperpolarization (Figure 5A). This strikingly precise spiking could be seen across PV interneurons, when pooling spikes across all cells and trials (Figure 5B). Interestingly, such brief odor-onset spiking was largely absent in SST cells (Figure 5C).

**Figure 5:**
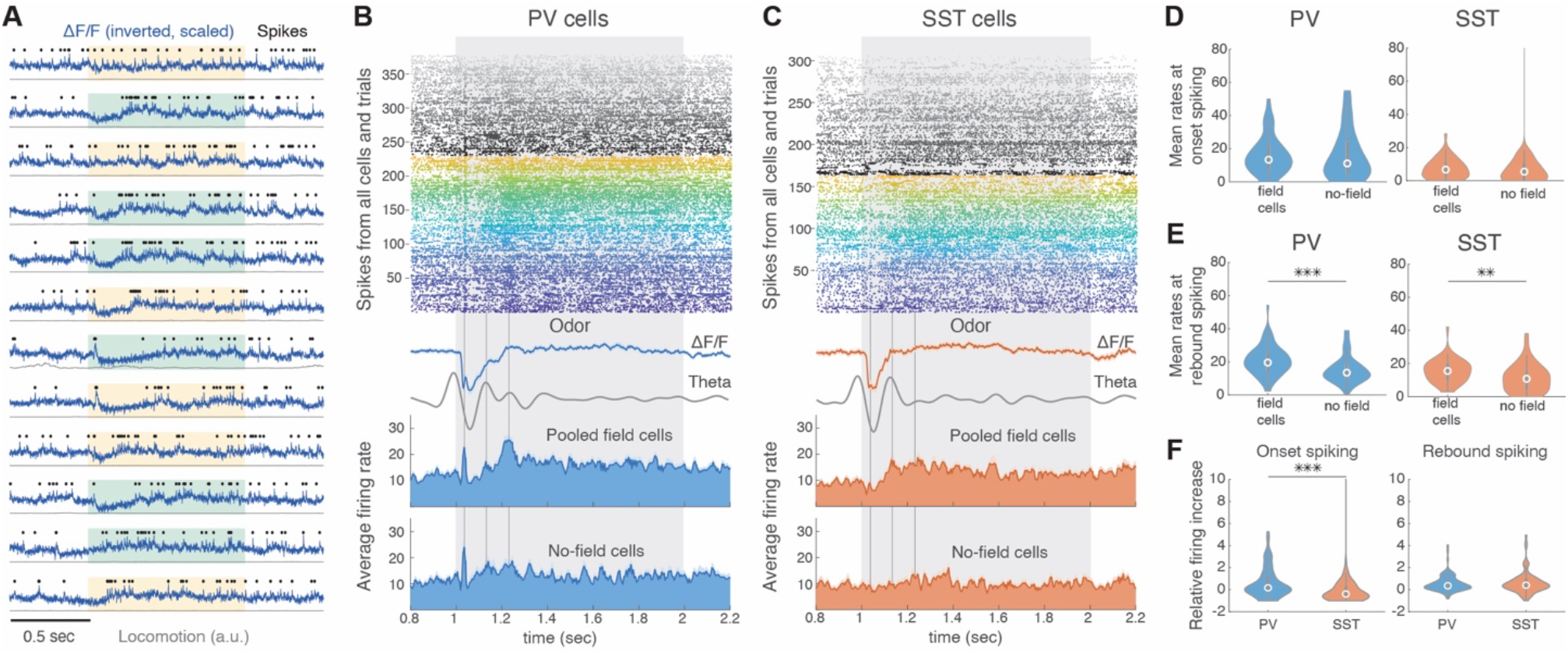
Brief synchronous spiking by PV cells, precedes PV and SST hyperpolarization and synchronizes rebound activity at theta peaks. **A**. Example trace of a PV cell around the first odor of a series of DNMS trials, displayed as before. Note the brief ‘onset spiking’, preceding the hyperpolarization in some, but not all, trials. **B**. Top: Spikes of pooled PV field cells (colored) and no-field cells (grayscale) around the first odor across all trials (each cell shown in separate color). Bottom: Average fine-scale firing rates (binned at 5 msec) of the two cell groups aligned to the mean ΔF/F of all PV cells (blue trace) and its theta-power bandpassed signal (gray). The three vertical lines are aligned to the first spike peak and the two theta peaks, respectively. **C**. Same for pooled SST cells. The three vertical lines indicate the same timepoints as in **B. D-E**. Average firing rates at onset spiking (30-50 msec after odor onset; **D**) and rebound spiking (200-500 msec after odor onset; **E**) in field cells vs no field cells. ** P < 0.01, *** P < 0.001, WT. **F**. Firing increase relative to baseline (0.5 sec pre-odor) at onset spiking (left) and rebound spiking (right) in PV vs SST cells. *** P < 0.01, WT. Single outlier truncated at **D** and **F** for plotting clarity.

To quantify these fast dynamics, we computed firing rates using 5 msec bins (unlike 100 msec before). Average firing rates across PV cells with and without a field, contained a ~30 msec peak at ~25msec after odor onset, superimposed on the average hyperpolarization. Moreover, the increase in odor-spiking after the hyperpolarization was not uniform throughout the remaining odor interval. Instead, average rebound spiking at the end of the hyperpolarization was briefly modulated by theta cycles, corresponding to the reset intracellular theta peaks (Figure 5B). The rebound spiking of SST cells was relatively attenuated, though a theta modulation was still present in field cells (Figure 5C). Since average intracellular theta was in phase between PV and SST cells (Figure 4K), the rebound spiking peaks were also synchronous between PV and SST field cells (Figures 5B-C). Similar features were observed between odor-A and odor-B trials (Figure S5) and across preferred and non-preferred odors in odor-specific cells (though, as expected, spiking was overall higher in preferred odors; Figure S5)

Odor-onset firing rates (30-50 msec post odor-onset) were similar between field and no-field cells (Figure 5D), but the rebound firing rates (200-500 msec post odor-onset) were significantly higher in PV and SST field cells compared to no-field cells (Figure 5E). As aforementioned, PV cells had a higher firing increase than SST cells during odor-onset, but the subsequent rebound firing increase was comparable between the two cell types (Figure 5F).

Finally, odor-onset spiking and rebound spiking in PV cells did not change between naïve and trained sessions. However, rebound spiking increased in SST cells after training (Figure S5).

Our findings reveal that the onset of hyperpolarization coincides with a brief synchronous PV spiking while its offset synchronizes rebound PV and SST spiking in theta peaks.

### Pyramidal odor responses are tuned by interneuronal spiking

The prevalence of interneuronal non-specific odor fields, implies that many pyramidal cells should be inhibited during odor delivery. We thus hypothesized that a large percentage of pyramidal cells would reduce their firing or become silenced during the odor.

To test this hypothesis, we analyzed a set of two-photon calcium imaging traces from dorsal CA1 of transgenic mice (N = 11), virally infected with synapsin-driven GCaMP6f, while mice performed the DNMS task [5]. Active pyramidal cells were segmented and GABAergic cells, expressing tdTomato, were excluded from analysis (Figure 6A). Odor-specific pyramidal sequences were previously observed in these recordings [5]. Here we used the deconvolved signal from pyramidal calcium traces, recorded at 30.9 frames/sec, as a proxy of spiking probability.

**Figure 6:**
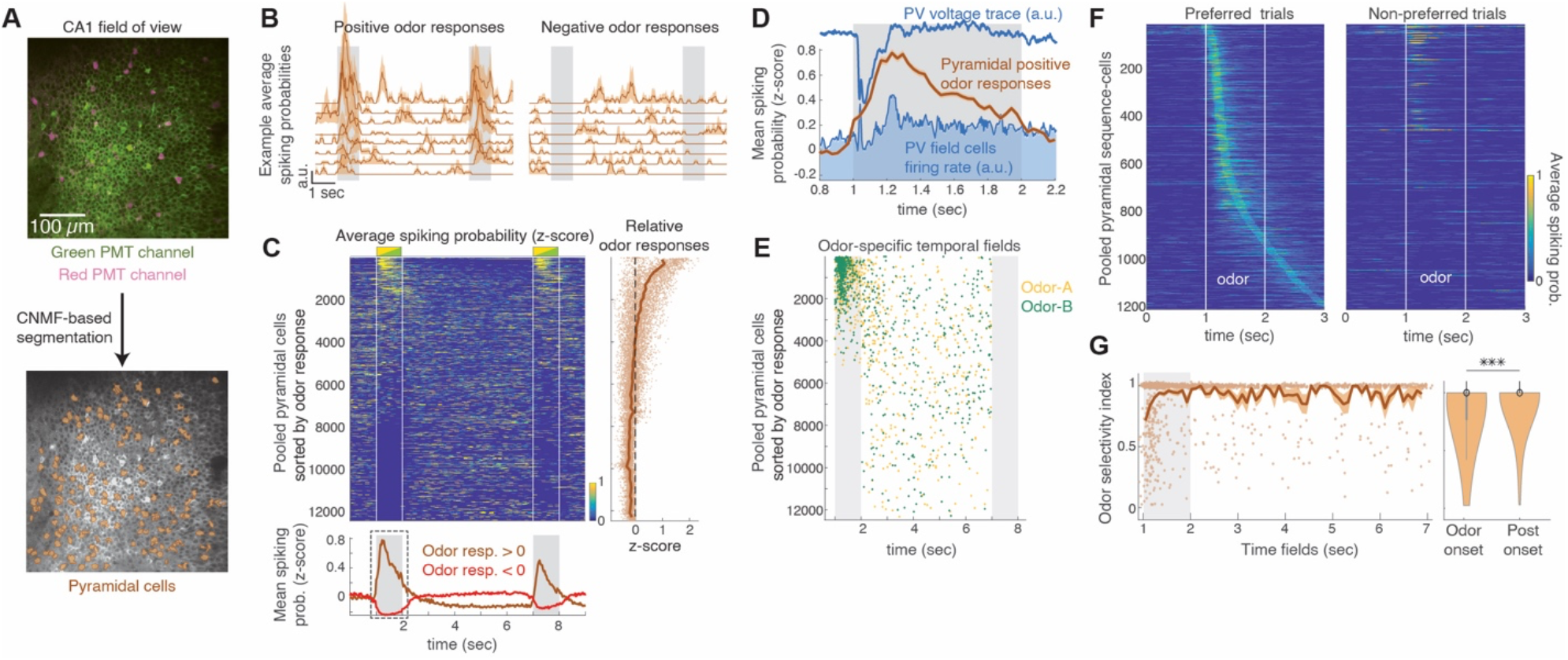
Pyramidal cell odor-responses are shaped by inhibition. **A**. Example field of view from two-photon calcium imaging in CA1 pyramidal layer of a Gad2Cre:Ai9 mouse expressing GCaMP6f (green) in all cells and tdTomato in all GABAergic cells (magenta). Right: Same FOV after ROI segmentation of pyramidal cells. **B**. Example traces of average deconvolved signal (‘spiking’) across all trials in cells with positive (left) and negative odor responses (right). **C**. Average z-scored spiking of pooled cells (N = 11 mice, 58 sessions), stacked by their average response during the first odor. Note that many cells stop spiking during the odor. Right: z-scored odor-responses averaged across the odor duration. Most cells exhibit a negative response. Bottom: Mean spiking across all cells with positive (black) and negative (red) odor responses. **D**. The same mean spiking, zoomed in around the first odor (dashed box in **C**). The average voltage trace from PV cells and the firing rates of PV field cells are superimposed (from Fig. 5B). Pyramidal odor responses are highly synchronous with the interneuronal rebound spiking. **E**. Field time bins of pooled Odor-A (yellow) and Odor-B (green) sequence-cells, sorted according to the order in **C**. Most time cells (70.8%) have negative odor responses. **F**. Average spiking (scaled) of pooled pyramidal sequence-cells, sorted by time field. Spiking is zoomed around the first odor in preferred (left) and non-preferred trials (right). Cells with fields after this time window are omitted. Note the response in non-preferred trials of early odor cells. **G**. Distribution of odor selectivity index from all sequence-cells (dots; small jitter added for plotting clarity) and average (black line). Multiple early odor cells have low selectivity index. Right: Selectivity index of early odor cells (field < 250 msec post odor-onset) vs the remaining sequence-cells. *** P < 0.001, left-tailed WT.

As expected, some pyramidal cells increased their firing during odor delivery (‘odor cells’) but others reduced their firing rates throughout the odor (Figure 6B). We sorted pooled pyramidal cells (N = 12,476 pyramidal cells from 58 imaging sessions) by their mean spiking probability during the first odor, across trials. A large proportion of neurons completely stopped spiking during odor delivery even though they were active before and after (Figure 6C). Indeed, relative odor responses, compared to baseline, were negative for most neurons (67.3% of all cells; Figure 6C), confirming the hypothesized widespread inhibition.

Interestingly, pyramidal cells with negative odor responses, had higher activity during the delay period, compared to cells with positive odor responses (Figure 6C), suggesting that they may constitute time cells (i.e. have delay time fields). This was confirmed by detecting significant fields based on the deconvolved signal (same criteria as in [5]) and sorting them in the same order as before. As expected, 96.7% of odor cells had positive relative responses to the odor. In contrast, most time cells had a negative odor response (70.8% time cells; Figure 6E), indicating that these neurons stop firing during the odor.

We next asked whether the hyperpolarization of PV and SST cells at odor-onset is shared by pyramidal cells. If it is shared, odor cells should spike after the hyperpolarization (~200msec) whereas if hyperpolarization is interneuron-specific, then odor cells should maximize spiking at odor-onset when they are disinhibited. We found that the average spiking probability of positive-responding pyramidal cells peaked between 200-400msec after odor-onset (Figure 6D). When superimposing on that activity the average voltage imaging trace and firing rate of PV cells, the peak in pyramidal activity was aligned with the peak in PV rebound spiking after their hyperpolarization, as well as with PV intracellular theta peaks (Figure 6D). This indicates that pyramidal responses coincide with interneuronal rebound spiking.

Finally, we asked if the coordination between pyramidal and interneuronal odor-spiking is important for pyramidal odor selectivity. Indeed, we found that pyramidal odor cells with an early field (in the first 250 ms after odor onset), during interneuronal hyperpolarization, tended to spike during both their preferred and non-preferred odor. (Figure 6F). As a result, average odor selectivity was significantly lower for fields within 250 msec from odor onset than later fields (Figure 6G).

Collectively, these findings confirm that increased PV and SST spiking during odor delivery inhibits most pyramidal cells and possibly allows them to have a field during the delay period. Moreover, there is a striking coordination between pyramidal and interneuronal rebound theta-modulated spiking which enhances pyramidal odor specificity.

## DISCUSSION

Hippocampal spiking sequences encode and retain sensory or contextual information across gaps in time [1, 2]. We investigated the role of interneurons in sculpting these sequences by employing fast rate voltage imaging *in vivo* of PV and SST interneurons in dorsal CA1. We used an odor-cued working memory task yielding odor-specific pyramidal sequences during odor delivery and the ensuing delay period [5]. Unlike pyramidal cells, interneurons predominantly encoded the timing of odor delivery but were mostly not odor selective, and few cells had time fields during the delay period. Odor-triggered inhibition was exerted by a stable number of cells with a constant flux of fields, irrespective of whether the animal was naïve to the task or trained to perform it. A transient and temporally precise PV burst at odor onset was followed by a widespread membrane potential hyperpolarization across many PV and SST cells. This hyperpolarization reset intracellular theta and organized rebound odor-spiking into theta-cycles synchronized across cells. Two-photon calcium imaging of pyramidal activity suggested that odor-timed inhibition silenced background pyramidal spiking, organized odor cell responses into a short time-window and enhanced pyramidal odor-selectivity. Therefore, PV and SST inhibition in CA1 increases the signal-to-noise ratio of odor cells during cue presentation, which is crucial for efficiently conveying sensory or contextual information downstream.

So far, most insights on the role of inhibition in hippocampal dynamics come from navigation tasks, often with contradictory results. For example, spatially tuned [32], untuned [33], and negatively tuned inhibition [34, 35] have been reported. Our findings reveal a combination of temporally tuned inhibition during cue presentation, followed by largely untuned spiking during the delay, suggesting that interneuron firing depends on the behavioral context. Interestingly, even though a minority of PV and SST cells reduced their spiking during odors (not shown), we did not systematically observe negatively tuned fields [34, 35].

Collectively, we found that inhibition sculpts sensory representations but less so internally generated ones (memory-encoding pyramidal time cells). This sculpting occurs in multiple ways:

1. Interneurons suppress background spiking during cue presentation through their widespread, largely non-selective activation. A similar background suppression role has been reported during spatial navigation, where untuned inhibition suppressed out-of-field excitation, enhancing the selective activation of place cells in each location [33]. In our recordings, this function was temporally limited to the cue presentation, yielding precise temporal tuning of interneurons.
2. By suppressing pyramidal activation during the odor, inhibition may allow time fields to emerge during the delay period. In spatial contexts, coordinated CA3 and entorhinal inputs [36] create dendritic plateau potentials on CA1 pyramidal cells, inducing near-by place fields through seconds-long behavioral-timescale synaptic plasticity (BTSP; [37]). Similarly, coordinated CA3-LEC inputs may induce plateau potentials on pyramidal cells during odor presentation. Without inhibition, most cells would burst and possibly obtain odor fields. Since BTSP extends for ~2 seconds after such event [37], followed by a ~4 second hyperpolarized membrane potential [38] (similar to the temporal structure of the odor-delay interval here), BTSP would enforce either an odor response or a delay timed-response, but not both. In that case, inhibitory suppression of odor-induced dendritic plateaus would be necessary to allow pyramidal cells to have an ensuing delay field. This is supported by our finding that most time cells were inhibited during the odor period. Dendritic inhibition by SST cells could be critical for such regulation.
3. Resetting of theta-paced interneuronal spiking organizes pyramidal odor responses into a narrow temporal window. PV cells can synchronize theta-paced rebound pyramidal spiking *in vitro* [39] and *in vivo* [17], mediated by hyperpolarization-activated inward current *I*_*h*_ [40], supporting our findings. A similar theta resetting was observed in EEGs from rat CA1 during odor sniffing, initiated at ~120-160msec after sniffing onset [4], similar to the timeline reported here. In the same study, odor-responsive pyramidal cells increased spiking ~300msec after sniffing onset, supporting the delay in pyramidal activity peak in our calcium data. Initial inhibitory responses of CA1 pyramidal cells, followed by resetting of theta-modulated spiking, were also observed in immobilized rabbits during sensory stimulation [41]. Restricting pyramidal responses temporally and organizing them into theta cycles, drives a higher entrainment of downstream cortical targets [42], efficiently transferring cue information. Further recordings with temporal resolution higher than calcium imaging are needed to pinpoint the exact temporal coordination between pyramidal cells and interneurons during odor presentation.
4. Interneurons promote pyramidal odor selectivity, since odor cells activated during interneuronal hyperpolarization have low selectivity. Assuming most pyramidal cells receive diverse LEC inputs onto their distal dendrites, possibly for either odor, cells would respond to both odors similarly. Odor-timed inhibition may sharpen odor selectivity by thresholding such responses, so that cells can spike only to the stronger odor-input. Analogous sharpening of tuning for visual orientation was reported to be driven by either PV cells [43, 44] or SST cells [45]. Given that most SST cells recorded here were possibly OLM cells, targeting distal dendrites where LEC inputs arrive, it is possible that they regulate this function. Future optogenetic SST- or PV-specific silencing during odor-onset will disentangle their role in shaping pyramidal odor selectivity.

Based on these observations, we predict that silencing either PV or SST interneurons during odor presentation would increase the number of odor cells (as seen with place cells in a similar condition [46]) but would decrease their odor-selectivity and the temporal precision in their responses and would reduce the number of time cells. This would significantly degrade both odor and time-information encoded by CA1 and possibly impair DNMS learning and performance.

The sparsity of interneuronal time fields during the delay period is in stark contrast with pyramidal time cell sequences. It suggests that inputs to time cells are either absent or not temporally tuned on PV and SST cells. Since most SST cells are driven by feedback CA1 excitation, they presumably receive inputs from multiple time cells with their time fields merging into continuous firing during the delay. The same may hold for PV cells though they receive inputs from CA3 as well, which also generates time cells [47]. We have previously argued that the relative stability of CA3 spatial maps, compared to the rapid remapping of CA1 spatial [48, 49] and time maps [5] makes CA3 less likely to be the sole source of CA1 time sequences during the delay. Time cells may instead be shaped by medial entorhinal sequences [50, 51], or even by LEC temporal coding [52]. Even though PV cells receive entorhinal inputs [53, 54], those do not trigger PV spiking [55]. Therefore, most PV responses are expected to be driven by Schaffer collaterals, supporting the theory that time cells depend on entorhinal inputs [56]. Additionally, theta-modulated medial septal inhibitory inputs on interneurons [57] may also cancel any temporally tuned excitation during the delay.

One of the most striking differences between PV and SST dynamics is the transient PV spiking at odor onset. Its ~20 msec temporal precision suggests that it is not generated by CA1 feedback excitation but is an externally driven response. Similar millisecond synchrony between pairs of interneurons has been observed during theta, gamma, and ripple oscillation [58]. Here, the synchrony extends throughout the PV population. PV cells are possibly responding to arriving excitation from CA3. The ~20msec between odor-onset and PV spiking agrees with the 15-25msec latency between olfactory bulb stimulation and hippocampal response in the rat [59, 60]. SST cells lack this response because they are mostly driven by CA1 PY recurrent excitation rather than CA3.

Multiple possibilities exist for the source of the widespread hyperpolarization: (1) It may be triggered by the odor-onset PV spiking, hyperpolarizing both interneurons and pyramidal cells simultaneously. This decrease in interneuronal activity, despite strong concurrent CA3 excitatory inputs, resembles paradoxical effects in inhibition-stabilized networks [61] and could be driven by strong PV recurrent inhibition [62]. (2) It may be driven by theta paced inhibition from the medial septum [57]. (3) It may be triggered by VIP cells which selectively target interneurons [63] and were recently shown to be recruited by LEC axons [64]. However, the disinhibitory effect of VIP cells boosts dendritic spike generation [64] and would probably result in increased pyramidal spiking during the hyperpolarization, instead of the other way around. (4) Direct long-range inhibition by LEC or MEC GABAergic cells could also produce a hyperpolarization on CA1 interneurons [65, 66], but not on pyramidal cells. The long axons, however, reach only interneurons in the stratum lacunosum moleculare, making this scenario less likely.

This work does not differentiate between different subtypes of PV or SST interneurons with distinct morphology and functional roles. It also does not include major classes like CCK and VIP cells. Future experiments are needed to disentangle the role of individual subtypes in spiking sequences. Moreover, recording from single cell-types prevents studying the interplay between the different cell types. Future recordings combining different GEVIs, expressed on different cell types will help address this problem [67].

Our findings highlight the power of voltage imaging in revealing cell-type-specific spiking and membrane potential dynamics during active behavior, with high temporal resolution [68, 69, 70] and across long time scales of several days. The recent developments of new GEVIs with improved fluorescence, kinetics and photostability [28, 30] render voltage imaging ideal for long-term recording membrane potentials from genetically identified cells *in vivo*. Future GEVIs will provide more photostability so that longer recordings can take place and CMOS technology development will allow larger fields of at higher rates so that neuronal populations can be recorded simultaneously. These advancements, together with the development of more targeted genetic markers, will soon allow delineating the role of specific interneuron classes in learning and memory behavior.

## ACKNOWLEDGMENTS

We would like to thank Stephen W. Evans, Alex Hao, Conor C. Dorian and Celina Yang for technical support. We are thankful to Sam McKenzie and Liron Sheintuch for their feedback. This work was supported by the US National Institutes of Health (R01 MH101198, RO1 NS099137, RO1 NS090930, R01 MH105427), the Intellectual and Developmental Disabilities Research Center at UCLA (U54 HD87101) and a Veterans Affairs Merit Review Award (1I01BX001524-01A1).

## AUTHOR CONTRIBUTIONS

J.T. and P.G. designed the experiments. M.Z.L. provided viral constructs. B.M. and J.T. constructed training and experimental equipment and conducted experiments. M.M. conducted experiments. J.T. analyzed experimental data. J.T and P.G. wrote the manuscript.

## METHODS

### Animals

A total of 10 adult male mice (12-21 weeks old) were used for *in vivo* voltage imaging experiments with the ASAP3 GEVI: 5 PV-Cre mice (B6;129P2-*Pvalb*^*tm1(cre)Arbr*^/J) and 5 SST-IRES-Cre mice (*Sst*^*tm2*.*1(cre)Zjh*^/J). For ASAP4 recordings, 5 adult SST-IRES-Cre adult mice were used.

For calcium imaging recordings, 11 adult mice (11-31 weeks old) were used: 5 Gad2-Cre:Ai9 mice (Gad2^tm2(cre)Zjh^/J crossed with B6.Cg-Gt(ROSA)26Sor^tm9(CAG-tdTomato)Hze^/J) and 6 Gad2-Cre:Ai14 (Gad2^tm2(cre)Zjh^/J crossed with B6.Cg-Gt(ROSA)26Sor^tm14(CAG-tdTomato)Hze^/J).

All animals were acquired from the Jackson Laboratory and were group housed (2-5 per cage) on a 12 h light/dark cycle. All experimental protocols were approved by the Chancellor’s Animal Research Committee of the University of California, Los Angeles, in accordance with NIH guidelines.

### Surgical procedures

Surgical procedures were described in detail by Taxidis et al. 2020 [5]. Briefly, isoflurane-anaesthetized mice were stereotactically injected with 500nl undiluted AAV8-ef1α-DiO-ASAP3-Kv (titer 3.83×10^12^ µg/ml), using a Nanoject II microinjector (Drummond Scientific) into the right dorsal CA1 area (−2mm posterior and 1.8 mm lateral to bregma, 1.3 mm ventral from dura) at 60 nl/min. 1 hour afterwards, a circular craniotomy (3mm diameter) was made around the injection site and the cortical tissue above the dorsal CA1 was aspirated until partial removal of the corpus callosum. A 3-mm titanium ring with a glass coverslip attached to its bottom was implanted into the aspirated area and secured to the skull. A custom-made lightweight metal head holder (headbar) was attached to the skull posterior to the implant. Cyanoacrylate glue and black dental cement (Ortho-Jet, Lang Dental) were used to seal and cover the exposed skull. Mice were administered carprofen (5 mg per kg of body weight) for 3 days post-surgery, as well as amoxicillin antibiotic (0.25 mg ml^−1^ in drinking water) through the water supply for 5 days.

For ASAP4 recordings, the same procedure was followed but 2 animals were injected with AAV8-ef1α-DiO-ASAP4b-Kv (titer 2.36×10^12^ vg/mL) and 3 animals were injected with AAV8-ef1α-DiO-ASAP4e-Kv (titer 3.45×10^11^ vg/mL). All ASAP viruses were produced by the Stanford Neuroscience Gene Vector and Virus Core facility.

For two-photon calcium imaging, the same procedure was followed but animals were injected with 1500 nl of 1:10 saline-diluted AAV1.Syn.GCaMP6f.WPRE.SV40 virus (diluted immediately prior to surgery; titre: 4.65×10^13^ GC/mL; Penn Vector Core).

### Experimental setup

The behavioral rig used for DNMS recordings was described in Taxidis et al. 2020 [5]. Briefly, mice were headfixed through a custom-made headbar holder on an 8-inch Styrofoam ball treadmill rotating in 1D. Locomotion was recorded using a custom printed circuit board based on a high sensitivity gaming mouse sensor (Avago ADNS-9500), connected to a microcontroller (Atmel Atmega328). A constant stream of clear air (~1 L/min) was supplied to the mouse through a custom-made lickport. During odor stimulation a dual synchronous 3-way valve (NResearch), switched from the clear air stream to the odorized air for 1 sec. Odorized air was created using a 4-ports olfactometer (Rev. 7c; Biology Electronics, Caltech), supplying air to either of two glass vials containing liquid isoamyl acetate (70% isoamyl acetate basis, FCC; Sigma Aldrich) or pinene ((−)-α-Pinene, ≥97%, FCC; Sigma Aldrich) odorants, diluted in mineral oil at 5% concentration. Odorized air reached the rig through Tygon tubing, leading to a dual 3-way solenoid valve (Lee Company). The olfactometer supplied air to either odor vial 1 sec prior to actual stimulation to allow for odorized air to travel through the tubing. During the 1 sec stimulation, the 3-way solenoid released it to the mouse through the lickport. At the offset of the stimulus, the valve switched the airstream back to clear air, ensuring a constant flow of air to the mouse and a quick clearing of the odorant from the air around the mouse. The odorized and clean air were set to similar airflow values (~1 lt/min), measured with a flowmeter (AWM3300; Honeywell). Licking was detected using a battery operated, custom-made, printed circuit board operating as a lickometer. Water droplets (~10 µl) were released by a solenoid valve (Lee Company). At the end of each trial, after the response window, vacuum was applied for 3 sec, to clear any lingering water and remove any odorized air. An additional 7 sec of intertrial interval were applied after the vacuum. The behavioral rig was controlled with custom written software (Matlab) and through a data acquisition board (USB-6341; National Instruments).

Voltage imaging was performed with a custom-built, high-speed, single-photon, epi-fluorescent microscope. Photoexcitation was provided via a fiber-coupled LED (Thorlabs, M455F3) with a center wavelength of 455nm. Excitation light was collimated after a 2-m long, 400-μm core multi-mode fiber optic patch cord (Thorlabs, M28L02) and expanded using a Keplerian telescope. The expanded beam was passed through a spectral excitation filter (Thorlabs, MF455-45), and reflected off of a long-pass dichroic mirror (Thorlabs, MD480) before reaching a 16×/0.8NA water-immersion objective (Nikon, CFI75 LWD 16X W). The expander was used to generate a localized excitatory spot ~165 μm in diameter, at the focal plane. Emitted fluorescence was collected and transmitted through the dichroic mirror and an emission filter (Thorlabs, MF530-43) before reaching a 100-mm tube lens (Thorlabs, AC300-100-A) to form an image on a fast scientific CMOS camera (Hamamatsu Photonics, ORCA-Lightning C12120-20P) capable of kHz framerates. The spatial sampling rate of the microscope was calculated as ~688nm and experimentally confirmed using a calibration test target.

Calcium imaging was described in Taxidis et al. 2020 [5]. Briefly, a resonant scanning two-photon microscope (Scientifica) was used, recording 512×512 pixel frames at 30.9 Hz, with the same objective as for voltage imaging (500×500 µm field of view). Excitation light was delivered with a Ti:sapphire excitation laser (Chameleon Ultra II, Coherent), operated at 920 nm. GCamp6f and td-Tomato fluorescence were recorded with green and red channel gallium arsenide photomultiplier tubes respectively (GaAsP PMTs; Hamamatsu). Microscope control and image acquisition were performed using LabView-based software (SciScan).

For all recordings, imaging and behavioral data were synchronized by recording TTL pulses, generated at the onset of each imaging frame, as well as olfactory stimulation digital signals at 1 kHz, using the WinEDR software (Strathclyde Electrophysiology Software).

### Behavioral training and voltage imaging protocol

Approximately, 2-4 weeks after surgery, mice were imaged for 2 sessions (separate days) while completely untrained in the DNMS task (one SST-Cre mouse was imaged for 3 sessions). They were presented with sets of DNMS trials and could passively smell the odor cues. Locomotion on the treadmill was typically limited since mice were not accustomed to the rig. After the second session, water restriction was initiated and DNMS training begun in separate training rigs. Protocols for water restriction and training to the DNMS task, starting ~10 days after surgery, were described in Taxidis et al. 2020 [5]. After mice reached performances > 90% for at least one 20-trial set (75% for one animal; one animal never reached criterion but only two cells were recorded post-training), voltage imaging resumed. The recording protocol was the same for naïve and trained performance (see below). Daily recordings typically stopped when either the mouse had stopped performing due to water satiation or no more cells with sufficient signal could be detected.

All imaging was performed under the custom-made single-photon microscope at 1000 frames per second. Before each recording, we searched for cells with strong epifluorescence through a 35×352 µm FOV (256×256 pixels and 2×2 pixel-binning). Once a cell was located, the FOV was reduced to 88×44 µm (64×32 pixels with 2×2 pixel-binning) which allowed for longer camera exposure (0.98 msec) and digitization at 1kHz frame rate. The FOV contained a single cell in most cases with only 3 PV-cell recordings (2.8% of all sessions) and 2 SST recordings (2.2%) containing >1 cells within the FOV with all cells yielding action potentials. Before DNMS recordings, 3 sec of activity were recorded. Using ImageJ, we manually segmented the recorded video and examined the raw fluorescence of the ROI. Only cells that yielded clear action potentials within the 3 sec video were further recorded. The power of the LED was adjusted depending on the signal-to-noise ratio in the 3 sec recording. LED power was typically set as low as possible to avoid fast photobleaching but also allow large spike amplitudes. The number of DNMS trials to be delivered was also set based on an assessment of the signal and the rate of photobleaching.

At each DNMS trial, the LED was on from 1 sec before the 1^st^ odor onset until the offset of the response window (11 sec total). It remained off during the 10 sec total intertrial interval. Frames were recorded only during the same time interval. After a set of trials was recorded, a time break of ~5-10min was given to prevent photo-toxicity. During this break, the acquired raw traces were examined, using ImageJ, for the existence of observable action potentials. If action potentials persisted throughout the final trial, a new recording of the cell was initiated over an additional set of trials. Otherwise, the cell was considered photobleached. In most cases, recording of a cell stopped after an adequate number of trials was recorded, before the cell was bleached. This allowed us to record cells for multiple consecutive days. After recording a particular cell, the FOV was expanded back to 352×352 µm and a small number of frames were recorded. The average frame of this video (ImageJ) was used as reference to locate the same cell on following days, based on cell-body shape as well as processes and surrounding features.

For experiments with odors turned off, the olfactometer ports were manually deactivated in every other trial, halting any flow of odorized air. Trials were otherwise normal, including all auditory cues from the 3-way valve. For detached air-valve experiments, the 3-way valve was manually detached from the rig for a subset of trials but remained connected and operational.

### Voltage imaging data processing

Each recorded video, together with the corresponding behavioral data, was processed separately using the Volpy automated pipeline for voltage imaging in Python [71]. The video was first motion corrected using the rigid NoRMCorre [72] algorithm as implemented in the CaImAn toolbox [73]. Parameters like the size of a kernel for high-pass spatial filtering or the maximum rigid shift were adjusted between videos to achieve efficient motion correction. ROI bounds were manually drawn around the recorded cell(s) using a graphical user interface in Volpy. Trace extraction and denoising and spike extraction took place reiteratively, based on the CNMF algorithm [74] and a modified version of the SpikePursuit algorithm [75], respectively. ΔF/F was computed as *(t-t*_*sub*_*)/t*_*sub*_ where *t* is the refined temporal fluorescence trace and *t*_*sub*_ is an estimated subthreshold trace. ΔF/F was inverted for all ASAP3 recordings but not for ASAP4 recordings. Spike detection was based on an adaptive threshold method. Parameters involved in trace denoising and spike detection, such as frequency for high-pass filtering to remove photobleaching or the maximum number of spikes to form a spike template, were adjusted between videos to ensure the accurate detection of action potentials that could be seen in the raw trace. Processing took place over the entire video with concatenated trials; traces and spike times were then split into the corresponding trials.

### Analysis of voltage imaging data

All analysis was performed using custom written software in MATLAB.

#### Motion Analysis

The locomotion signal was recorded at 1 kHz. Its mode value was subtracted, it was turned to absolute values and was smoothed with a 1^st^ order Savitzky-Golay filter. Motion segments were taken to be those were the locomotion signal exceeded 0.02 and immobility was assumed when it dropped 0.01 (a.u.). Consecutive motion segments closer than 20 msec were concatenated and segments shorter than 10 msec were discarded.

#### Spiking and subthreshold membrane potential analysis

All processed sets of trials from each neuron in a session, along with the behavioral data, were concatenated and considered one continuous recording of that cell. The quality of processing from each recording was assessed visually. Cells that yielded signals that were too noisy, with problematic spike detection or with no spikes, were removed from analysis. Trials at the end of a recording containing very few or no spikes, where the cell was photo-bleached, were also removed.

Firing rates were computed by binning the extracted spike times of each trial at 100 msec or 5 msec (for Figure 5 and S5) non-overlapping bins. Inter-spike intervals were computed for each trial separately. The burst index of a neuron was quantified as the percentage of spikes closer than 10 msec, over its total number of spikes.

Speed scores were computed as the trial-averaged correlations between a cell’s firing rates (smoothed with a 5-point moving average window in each trial) and the animal’s locomotion in the corresponding trials (downsampled to 1 kHz sampling rate to match the firing rate sampling rate). Odor scores were the trial-averaged correlations between a cell’s firing rate and a vector of trial length, containing 1s in the odor-delivery timebins and 0s in all other timebins.

Frequency-domain analyses of subthreshold membrane potentials were based on despiked versions of the ΔF/F signal. Spike removal was performed through a Bayesian method, developed for removing spike waveforms from local fields potentials (LFP) to eliminate artifactual spike-LFP correlations [76]. A segment of 25 msec around each detected spike (5msec before and 19msec after the action potential peak) was replaced in the ΔF/F based on Bayesian inference. Due to the action potential widening by GEVIs, this segment encompassed the average recorded action potential and the ensuing after-hyperpolarization.

Power spectra were computed separately on concatenated motion or immobility despiked ΔF/F segments from each cell, through a modified periodogram using a Kaiser window with a frequency resolution of 0.5 Hz. Delta and theta phases and amplitudes were computed from the Hilbert transform of the despiked ΔF/F after bandpassing it in the 0.5-3Hz or 4-10Hz frequency range, respectively, using a 3^rd^ order Butterworth filter. The phase of each spike was computed as the angle of the Hilbert transform at the spike peak. Delta and theta amplitude traces were smoothed with a 1^st^ order Savitzky-Golay filter. Average amplitudes were estimated as the mean amplitude across all trials and all timepoints. Average theta phase of a cell’s spikes and mean vector length (strength of theta modulation) and phase variance were computed using the Circstat toolbox [77].

For hyperpolarization analysis, non-despiked ΔF/F traces were smoothed by 1^st^ order Savitzky-Golay filter and z-scored to the mean and standard deviation of the 0.5 sec preceding the onset of the first odor (baseline) in each trial separately. Time segments where this z-scored trace *S* was *S* < −1 during to the first odor delivery were detected. The first continuous time segment *T*_*h*_ (including all segments closer than 20msec) was considered a potential hyperpolarization. If *T*_*h*_ had total duration of > 50 msec and contained timepoints where *S* < −3 with total duration >10msec, then *T*_*h*_ was considered a significant hyperpolarization. The first timepoint, the total length and the *S* minimum value of *T*_*h*_ were taken as the hyperpolarization onset time, duration, and amplitude respectively. The timing of minimum *S* value was taken as the hyperpolarization trough timing. For comparisons including low number of recorded cells (Fig. S5), the minimum value of *S* within the first 200 msec after odor onset, in every trial, was taken as the hyperpolarization amplitude.

For fine-timescale analysis (Fig. 5 and S5), firing rates (5 msec bins) were smoothed with a 3-point moving average window. Odor-onset and rebound spiking corresponded to the bins within 30-50 msec and 200-500 msec after odor onset, respectively. Corresponding average rates referred to the mean firing rate value across the respective time bins and across all trials. Firing increase was computed as the relative change of the trial-averaged rate, averaged across the corresponding timebins, to its baseline average rate (average over the 0.5 sec preceding the odor).

All plotted ΔF/F traces were normalized by their maximum value over each trial. Plotted firing rates over individual trials, as well as average rates, were z-scored based on their mean and SD across the odor-delivery interval. Firing rates or theta amplitude traces were smoothed by a 5-point moving average window for plotting.

#### Field detection and analysis

The firing rate across each trial was smoothed with a 5-point moving average window. Firing rate signals over the odor-delay interval (from onset of first odor to end of the delay period) were z-scored in each trial and split over trials initiated by odor A or odor B. The average rate was computed for each set and the corresponding maximum values were noted for each. The two sets of firing rate traces were then circularly shifted by a random interval up to ± 1/2 odor-delay interval duration, separately for each trial, and the maximum average firing rate over the shifted trials was again computed. This process was repeated 1000 times, generating a distribution of maximal rate values. The cell was considered to have a significant firing field for a particular odor, if the corresponding maximum mean firing rate was larger than the 95th percentile of the shuffled distribution. The time-bin of that maximal rate was considered the cell’s field time-bin. Cells with firing fields for both initiating odors were considered non-odor-specific. For these cells, if their field time-bins for each odor were closer than 1 sec, then their final field was considered the average of the two time-bins. If their field time-bins were further apart than 1 sec, then their field was the bin where the maximal average rate was higher. This only occurred in 3 PV cells (2.8% of all PV cells) and 4 SST cells (4.3%). For cells with no field, the time-bin of their maximal average firing rate over all trials was computed.

The selectivity index of each cell was then computed as:

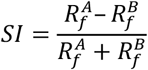

where 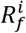 is the cell’s average firing rate at its field (or at the time-bin of maximal average firing rate in the case of no-field cells) over all trials initiated by odor *i*. Cells that were initially detected as being odor-specific but had |SI| < 0.42 were considered as non-odor-specific. This criterion was implemented because many cells with similar firing rates in both odors were erroneously classified as odor-specific due to noisy spiking in other time-bins as well.

A cell’s firing rate change at different trial segments was computed as the trial-averaged relative change of its mean rate over the corresponding time-bins, compared to its mean rate over the 0.8 sec preceding the first odor (baseline).

#### Analysis Across Multiple Days

The inflow of new odor fields at any session was computed as the percentage of cells of day *d* with an odor field that were also imaged in the previous session but did not exhibit a significant odor field then. Stable cells are defined as the percentage of cells of day *d* with an odor field that were also imaged in the previous session and did exhibit a significant odor field then too. The outflow of odor fields at a session was computed as the percentage of cells from the previous session with an odor field that were also imaged on the current one but did not retain an odor field.

For comparing between the firing rates or theta oscillations of tracked cells while the mouse was naïve vs trained, we used the first trained session where the cell was imaged to compare against the one naïve session where the cell was imaged (for one PV cell which was imaged on two naïve sessions, we used the second session).

For cross-days correlations, firing rates over the odor-delay interval were z-scored for each trial and the Pearson correlation was computed for all pairs of trials between the two sessions. The average correlation was computed. The Spearman rank correlation was estimated for the relationship between mean pairwise correlations and distance between imaging sessions.

When estimating the evolution of collective activity from all cells across days, the 1 extra naïve session in a single SST-Cre mouse was removed for analysis (so that all mice had 2 naïve sessions). Performance was computed over the pooled trials from all trial blocks recorded with each cell. Rates, locomotion, and theta amplitudes were averaged across the entire recording. For naïve sessions all performances were set to 50% corresponding to chance, since mice did not participate to the DNMS task.

Theta power refers to the average theta amplitude over odor-delay interval and across trials.

#### Support Vector Machine and Bayesian Decoding

For both decoding methods, trials from each cell were analyzed separately. The first 2/3 of all trials were used for training and the final 1/3 for decoding. Only recordings with at least 10 trials total and at least 2 training trials of each odor were included. Rates over each trial were smoothed with a 5-point moving average window.

Binary support vector machine (SVM) classifiers were created, to decode the identity of the first odor in a trial. They were trained on the cell’s firing rates either across the first odor delivery or the entire odor-delay interval. Smoothed rates were z-scored over the corresponding bins in each trial. The classifier was trained using a radial basis function kernel with automatically selected scale factor and standardized predictors (function *fitcsvm*, MATLAB). Chance baseline for each cell was computed by shuffling the identity of both training and predicted odors with 500 repetitions and applying the SVM classifier on each repetition. Odor-decoding accuracy refers to the percentage of correctly decoded odors over all predicted trials.

Bayesian decoders were created to decode odor-specific timing in a trial, using a similar approach as in Taxidis et al. 2020 [5]. They were trained on the cell’s firing rates across the odor-delay interval. The rate of each trial was scaled to have a maximum value equal to the maximum over all trials. To decode the odor identity, we considered time space to be 2 x odor-delay interval, by concatenating along the time axis, each cell’s mean rate over the two initiating odors (mean rate over odor-A-initiated trials followed by mean rate over odor-B-initiated trials). The decoder, trained by these concatenated firing rates, would thus predict a time point along a doubled time interval, 12 sec long. If the time point was within the first half (0-6 sec) it corresponded to that time point of an odor-A trial, whereas if it was within the second half (6-12 sec) it corresponded to the analogous time point of an odor-B trial. Time bins with no activity in either group of trials were discarded.

The decoded time point 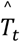 from the cell’s spiking at time-bin *t* in a decoded trial is given by:

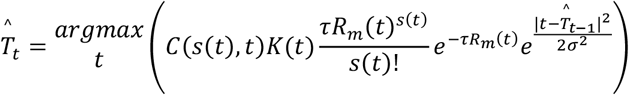

where *s(t)* is the number of spikes at time-bin *t, R*_*m*_*(t)* is the cell’s mean firing rate across odor-A trials concatenated with that across odor-B trials, at time-bin *t, τ* is the bin duration (100 msec) and *K* is the probability of being at time-bin *t* of a trial initiated by a particular odor (it is proportional to the ratio of trials of that odor over all training trials) and *C(t)* is a probability normalization factor. The last term functions as a continuity constraint, limiting the decoded time-bin to a relative proximity to the previous one, with *σ* = 3 sec. Chance baselines were computed by randomly circularly shifting the time-bins of each trial to be decoded, with 500 repetitions, and decoding the shifted trials for each repetition. Decoded time error refers to the mean absolute time distance between a given timepoint and the decoded one: 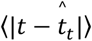, irrespective of the initiating odor. It thus functions as a measure of the time-information carried by the cell. Odor decoding accuracy refers to the percentage of correct odor decoding at any given timepoint (i.e., decoded time-bin being in the correct half of extended time-axis).

#### ON OFF trials

Locomotion and DFFs were smoothed with 1^st^ order Savitzky-Golay filter. Rates smoothed with a 5-point moving average window. For each cell, all 3 measures were normalized by their corresponding maximum value across all trials of the average value across the odor-delay interval.

### Two-photon calcium imaging and data processing and analysis

The recording protocol and calcium data processing was described in Taxidis et al 2020 [5]. Briefly, two-photon imaging sessions started ~2-3 weeks post-surgery while mice were either in DNMS training or were well-trained. A single fixed FOV was imaged every day for each mouse. A total of 58 recording sessions from 11 mice were included here. Calcium data were processed in MATLAB using a custom-built pipeline based on the CaImAn package [73]. GABAergic neurons were identified based on their static Td-Tomato fluorescence recorded for 500 frames at the beginning of each session on the red-channel PMT, together with the functional (green) channel. The ROI from each channel were then registered and ROI from the green channel that matched those from the red were discarded, so that only pyramidal cell activity from the green channel was analyzed further.

Pooled cells were sorted based on the mean value of the trial-averaged deconvolved signal, over the first odor in each trial. The trial-averaged deconvolved signal was then z-scored. The mean value of the z-scored signal over the first odor was used as the cell’s odor response. Detection of significant fields followed the algorithm in Taxidis et al. 2020 [5] but was applied to the raw deconvolved signal here, instead of its binned version.

When plotting pooled field-cell spiking (Fig. 6F), average deconvolved signals over preferred or non-preferred trials were normalized by the cell’s average signal at its field. Example deconvolved signals (Fig. 6B) as well as trial-average ones were smoothed with a 5-point moving average window for plotting.

