## Supplemental figures for "Voltage imaging reveals that hippocampal interneurons tune memory-encoding pyramidal sequences"

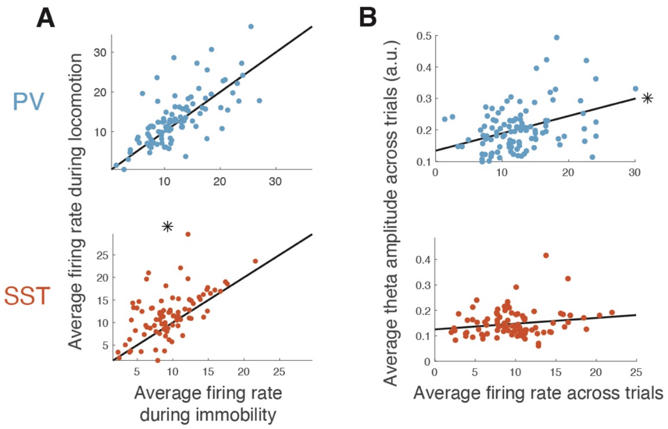


**Figure S1: Relationship between spiking, intracellular theta and locomotion in PV and SST cells. A.** Average firing rates for all PV and SST cells (dots) during locomotion versus immobility. * $P < 0.05$, paired-sample t-test. **B.** Correlation between average intracellular theta power and average firing rates is significant for PV cells only. * $P < 0.05$; random permutation test. Black lines: Least-squares fit.

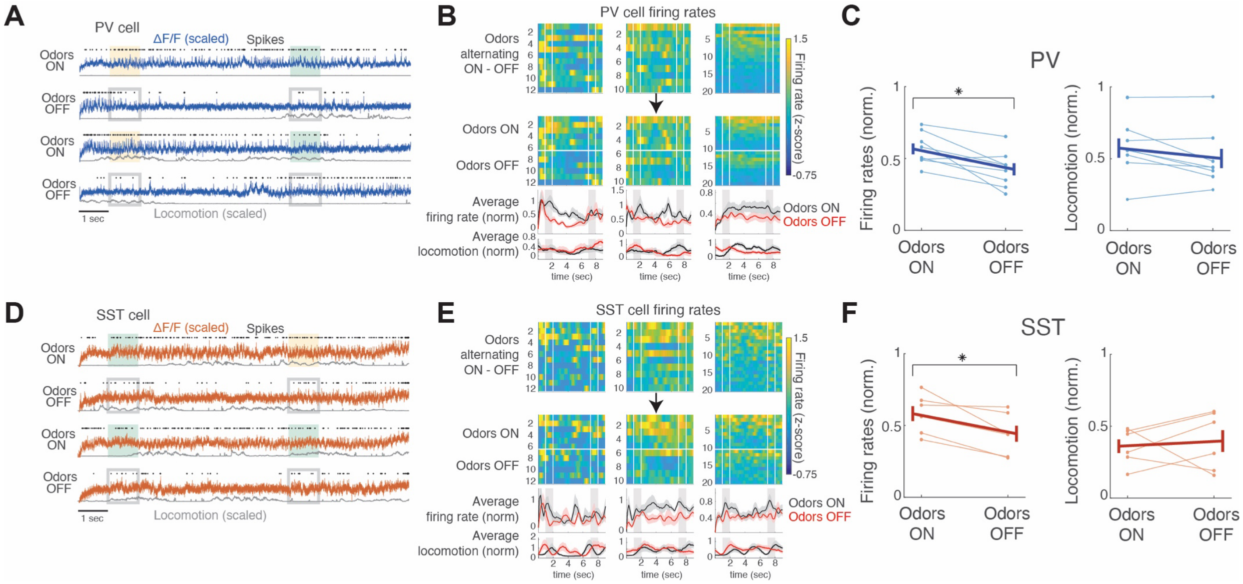


**Figure S2: Firing rates are lower when odors are turned off, but locomotion remains similar. A.** Example traces from a PV cell across 4 trials, displayed as in Fig 1C. Gray boxes: Odor cue windows in trials where the odor was not delivered. **B.** Top row: Firing rates from 3 PV cells across trials with alternating odors ON and OFF (scaled by maximum average rate across odor/delay bins). Second row: Same firing rates rearranged over all odor ON and OFF trials (separated by white line). Third row: Mean ± SE firing rates across odor ON and OFF trials (black and red respectively). Bottom row: Average locomotion in the same trials. **C.** Mean ± SE firing rates (averaged across all odor/delay bins and all corresponding trials) of each cell (left) and corresponding average locomotion (right) in odor ON versus odor OFF trials. * $P < 0.05$, paired-sample t-test. No significant differences exist for locomotion. **D-F.** Same as A-C for SST cells.

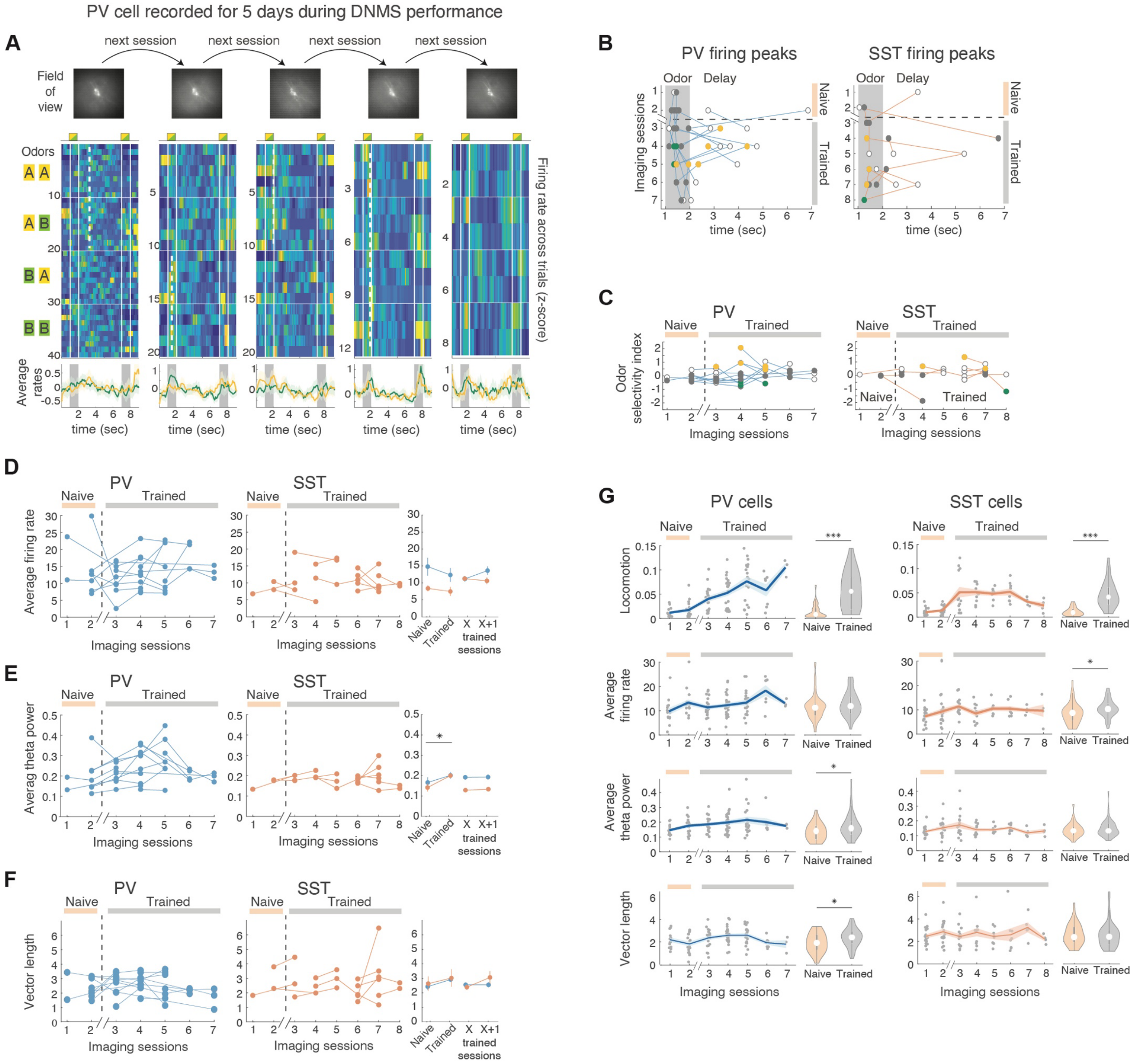


**Figure S3: Interneuron firing properties remain overall stable across days and over training. A.** Example PV cell recorded for 5 consecutive imaging sessions, plotted as in Fig. 3. Dashed lines indicate a significant field over a specific odor (dashed line covers corresponding trials) or through both odors (Day 4). **B.** Progression of firing peak time-bins across days for PV (left) and SST cells (right) recorded across multiple sessions. Grey: Non-odor-specific fields. Yellow, green: Odor-A- or Odor-B-specific fields, respectively. Open circles: non-significant peaks. Lines connect a cell's progression. **C.** Progression of odor selectivity index across days, displayed as in **B** (sessions on x-axis). **D-F**. Progression of average firing rate (**D**), theta power (**E**) and phase locking strength (mean vector length; **F**) across all trials, as in **C** (field-type not displayed). Right: Mean ± SE for last naïve vs first trained session and for every X vs X+1 trained session. * $P < 0.05$, paired sample t-test (distributions of PV and SST cells were pooled for this comparison. $P > 0.05$ for all other comparisons). **G.** From top: Evolution of average locomotion per session, mean firing rate per cell over all trials, theta power and theta modulation of spiking (vector length) per cell across PV-Cre mice and PV cells (left) or SST-Cre mice and SST cells (right), plotted as in Fig. 3I-J. * $P < 0.05$, *** $P < 0.001$, WT.

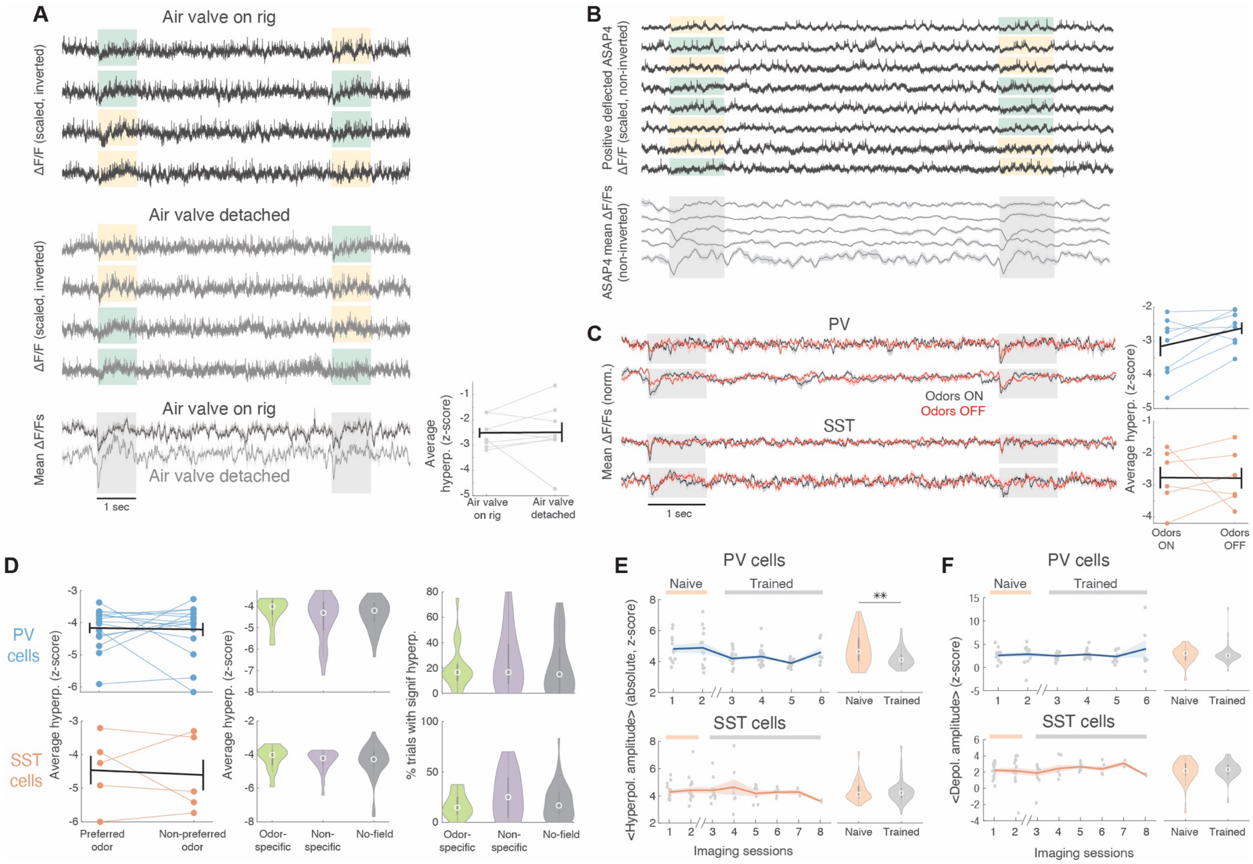


**Figure S4: Odor-onset hyperpolarization is not artifactual and is reduced in PV cells after training. A.** Example traces across 4 DNMS trials from PV cell exhibiting odor-onset hyperpolarization, when the odor air-valve is on the rig (top) versus detached (middle). Average ± SE ΔF/F over trials with the valve on the rig versus detached. Right. Average hyperpolarization (ΔF/F minimum, z-score-scaled over baseline of 0.5 sec before odor onset) in the two conditions ($P > 0.05$, WT). **B.** Top: Example traces from a cell expressing the ASAP4 GEVI (top) and average traces from 5 cells. ASAP4 is positively deflected with depolarization and traces are therefore not inverted. The odor-onset deflection is still indicating a hyperpolarization. **C.** Average traces from two example PV (top) and two example SST cells (bottom) across trials with the odors ON (black) vs OFF (red). Right: Average hyperpolarization (scaled as before) for the two cell groups over the two conditions. * $P < 0.05$, paired-sample t-test. **D.** Left: Average hyperpolarization over preferred vs non-preferred trials for PV and SST odor-specific field-cells. Middle: Average hyperpolarization in odor-specific, non-specific field-cells and no-field cells. Right: frequency of hyperpolarization occurrence for the three cell groups. **E.** Evolution of mean ± SE hyperpolarization across sessions and naïve vs trained session averages (right) for PV cells, displayed as in Fig. 3I-J, * $P < 0.05$ WT. **F:** Same for SST cells.

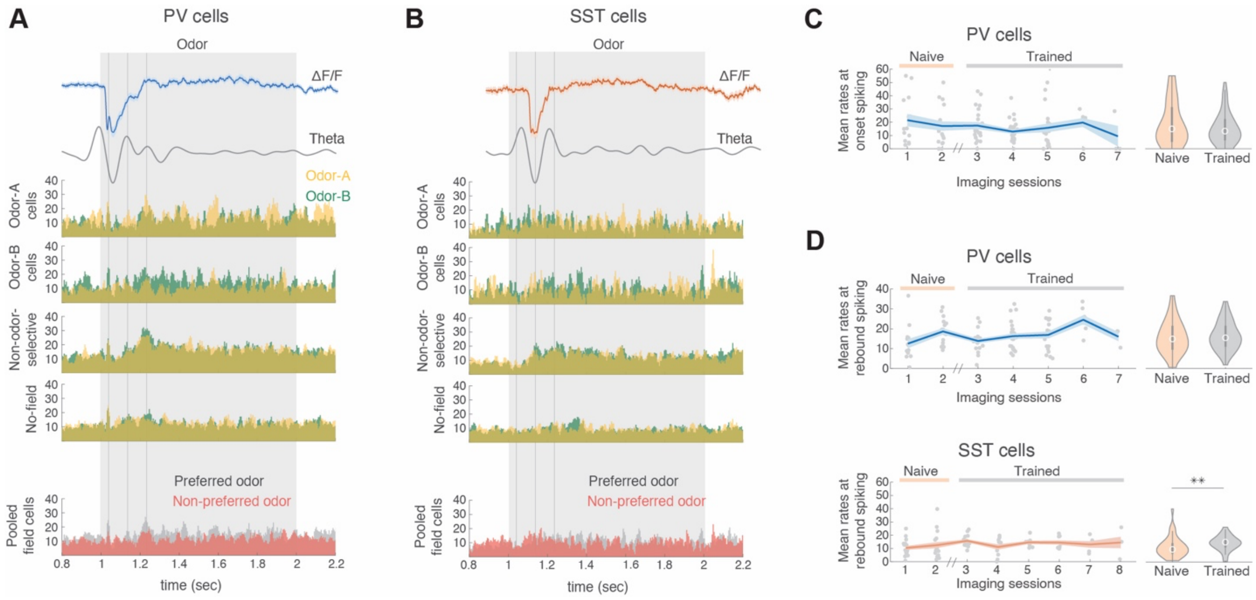


**Figure S5: Fine-timescale spiking across different cell groups and odors, and its evolution across days. A-B.** Average wideband and theta bandpassed ΔF/F aligned with mean firing rates of (from top) odor-A, odor-B, non-odor-specific and no field PV cells (**A**) and SST cells (**B**) across odor-A (yellow) and odor-B (green) trials. Bottom: Rates of odor-specific cells in their preferred (gray) and non-preferred trials. **C.** Evolution of odor-onset spiking in PV cells (SST cells did not exhibit onset spiking) plotted as before. **D.** Evolution of rebound spiking in PV and SST cells. ** $P < 0.01$ WT.
